# Engineering age-adaptive mRNA lipid nanoparticle cancer vaccines via reprogramming systemic gene expression

**DOI:** 10.64898/2026.04.11.717910

**Authors:** Yining Zhu, Christine Wei, Jingyao Ma, Di Yu, Jialiang Wang, Xiaoya Lu, Kailei D. Goodier, Jinghan Lin, Xiang Liu, Yunhe Su, Zhuoqing Jiang, Autumn H. Greco, Leonardo Cheng, Wu Han Toh, Yang Miao, Jonathan P. Schneck, Joshua C. Doloff, John W. Hickey, Hai-Quan Mao

## Abstract

Lipid nanoparticle (LNP)–based mRNA vaccines have transformed cancer immunotherapy, yet their efficacy in older individuals, who represent the majority of cancer patients, remains poorly understood. Here, we uncover a critical and previously underappreciated barrier to mRNA LNP cancer vaccine performance in aged hosts: impaired systemic transgene expression. Using the SM-102 mRNA LNPs as a benchmark formulation, we show that while local immune activation and antigen presentation at the injection site and draining lymph nodes remain largely intact with age, transgene expression in peripheral organs, including the liver, lungs, and spleen, is markedly reduced. This deficit limits the magnitude and durability of CD8 and CD4 T cell responses and substantially compromises tumour control. Transcriptomic profiling further reveals that attenuated transgene expression parallels broad attenuation of antigen processing, presentation, and activation pathways in immune cells from aged animals, implicating impaired systemic mRNA translation as a central driver of the downstream immune defects and antitumour efficacy loss. Building on these insights, we identify a rationally selected LNP formulation that reestablishes distal antigen expression across age groups as an engineering strategy to revive T cell immunity, achieving full rescue of therapeutic efficacy in aged mice without additional intervention. Together, these findings establish systemic mRNA translation as a tuneable lever of vaccine performance and highlight that optimizing LNP formulations to sustain systemic transgene expression across age groups may enable next-generation, age-adaptive mRNA vaccines for cancer and other diseases of aging.

## INTRODUCTION

Messenger RNA (mRNA) vaccines formulated with lipid nanoparticles (LNPs) have emerged as a transformative platform for cancer immunotherapy.^1–6^ Their safety, scalable manufacturing capabilities, and ability to elicit both humoral and cellular immune responses have accelerated clinical translation, with over one hundred cancer vaccine trials now in progress, targeting an array of tumour types including melanoma, non small cell lung cancer, colorectal, pancreatic, and more.^7–9^ In particular, Moderna’s mRNA-4157/V940 vaccine demonstrated improved recurrence-free survival in melanoma patients when combined with anti-PD-1 therapy.^10^ Early phase studies have reported favourable safety and immune activation profiles across indications, such as neoantigen-targeted vaccines for gastrointestinal and pancreatic cancers, with evidence of antigen-specific T cell responses in over half of vaccinated patients.^10–13^ These achievements have significantly accelerated the deployment of LNP-based mRNA vaccines.

LNP-based vaccines rely on a sequence of tightly coupled events: nanoparticle penetration in the tissue, cellular uptake, endosomal escape, cytosolic mRNA translation, and antigen processing and presentation to activate adaptive immunity.^3,13–15^ Formulation parameters, such as lipid chemistry, PEG-lipid content, and helper lipid identity, markedly influence these processes and thereby determine the magnitude and quality of immune responses, both local and systemic outcomes.^16–24^ While efforts to enhance potency have focused on optimizing lipid composition and targeting immune cells, most preclinical studies have used young adult models, leaving the impact of aging on mRNA vaccine performance largely uncharacterized.^25–29^

Aging is accompanied by immunosenescence, a decline in both innate and adaptive immune functions, reflected by diminished antigen presentation, reduced T cell priming, and weakened effector responses, as well as systemic alterations in metabolism and protein synthesis, collectively termed translational senescence.^30,31^ These changes influence endosomal processing, mRNA stability, and ribosomal efficiency, all of which could limit mRNA processing and translation in aged tissues. Currently, it remains unclear which stages of mRNA vaccine processing are most vulnerable to aging, and whether these effects arise from reductions in nanoparticle trafficking, biodistribution, immune sensing, or intracellular translation. Addressing these questions is critical, as the majority of cancer patients and vaccine recipients are older adults whose responses are unpredictable using current LNP designs.

In this study, we set out to systematically examine how aging influences the performance of mRNA-LNP vaccines, using the clinically benchmarked SM-102 LNP formulation as a model. By comparing young and aged mice, we sought to determine whether age alters LNP biodistribution, mRNA translation, antigen presentation, or immune activation, and to identify which stages of the vaccine response are most vulnerable to aging. We combined *in vivo* imaging, reporter mouse models, and transcriptomic profiling to dissect these processes at both local and systemic levels. This approach was further extended to evaluate whether formulation engineering could mitigate age-associated functional deficits. Together, these studies aim to reveal how aging reshapes the biological interface between mRNA translation and immune activation, providing mechanistic insight to guide the design of more effective mRNA vaccines for older populations.

## RESULTS

### Aging attenuates humoral and cellular immune responses to SM-102 mRNA LNP vaccine

To establish a baseline for how aging influences vaccine performance, we compared the immune responses of young (6–8 weeks) and aged (10–12 months) C57BL/6 mice following intramuscular (i.m.) administration of SM-102 mRNA-LNPs encoding ovalbumin (mOVA). Mice received three doses of 10 μg mOVA SM-102 LNP via i.m. injection on days 0, 7, and 14 (**Fig. 1a**). Serum antibody analysis on day 30 revealed markedly reduced titers of anti-OVA IgG, IgG1, and IgG2c in aged mice compared to young counterparts (**Fig. 1b–d**). IgG titers in aged mice declined to ∼21% of those in young mice; IgG1 and IgG2c levels were reduced to ∼34% and ∼25%, respectively, indicating a substantial reduction in humoral immunity.

**Figure 1.**
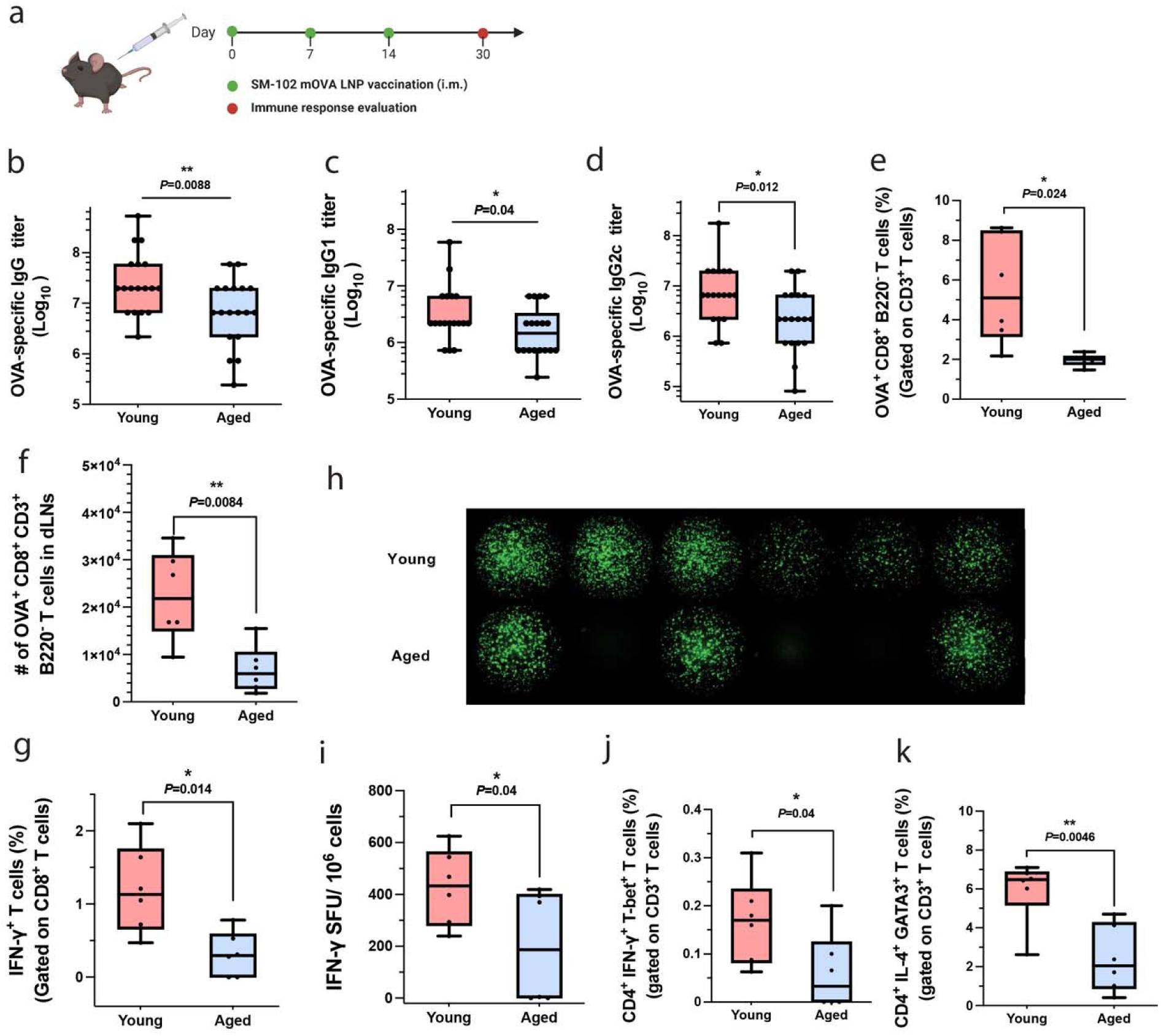
Aged mice exhibit impaired humoral and cellular immune responses following SM-102 mRNA-LNP vaccine. **a,** Schematic of the vaccination schedule. Young (6–8 weeks) and aged (10–12 months) C57BL/6 mice were vaccinated intramuscularly with 10 μg of mOVA-loaded SM-102 LNPs on days 0, 7, and 14, and sacrificed on day 30 for immune profiling. **b**–**d,** Serum antibody titres against OVA were measured by ELISA on day 30 post-vaccination, including total IgG (**b**), IgG1 (**c**), and IgG2c (**d**). **e**–**f,** Antigen-specific CD8⁺ T cells (CD3⁺B220⁻CD8⁺OVA⁺) in the draining lymph nodes (dLNs) were quantified by flow cytometry, shown as percentage among total live cells (**e**) and total cell number (**f**). **g**–**i,** Splenocytes were restimulated *ex vivo* with OVA (100LJμgLJmL⁻¹) and SIINFEKL peptide (2LJμgLJmL⁻¹) for 12 h. Cytokine-producing CD8⁺ T cells were analysed by intracellular cytokine staining for IFN-γ (**g**) and quantified by FluoroSpot (**h**–**i**). **j**–**k,** Percentages of CD4⁺ T helper cell subsets in spleens post-restimulation, including Th1-like cells (CD3⁺CD4⁺Tbet⁺IFN-γ⁺; **j**) and Th2-like cells (CD3⁺CD4⁺GATA3⁺IL-4⁺; **k**), were measured by flow cytometry. Data represent the mean ± s.e.m. (n = 18 mice per group for **b**–**d**; n = 6 mice per group for **e**–**k**). Data were analysed using an unpaired t-test. \**p* < 0.05, ** *p* < 0.01. Panel **a** was created in BioRender.

We next assessed antigen-specific T-cell responses. The frequency of OVA-specific CD8 T cells (CD3 B220 CD8 OVA) in the draining lymph nodes (dLNs) was reduced by 2.78-fold in aged mice (**Fig. 1e, f, Supplementary Fig. 1**), and the total number of OVA-specific CD8 T cells was similarly reduced (∼3.25-fold; *p* = 0.0084). In the spleen, cytotoxic T cell responses were evaluated by intracellular cytokine staining for IFN-γ in CD8 T cells. Aged mice exhibited a ∼3.78-fold lower percentage of IFN-γ CD8 T cells relative to young mice (**Fig. 1g, Supplementary Figs. 2** and **3**). These results were corroborated by IFN-γ FluoroSpot assays, which revealed a ∼2.16-fold higher frequency of IFN-γ–secreting cells in young animals (**Fig. 1h, i**). We further examined CD4 T cell polarization. The proportion of Th1 cells (CD3 CD4 IFN-γ Tbet) in the spleen was around 2.76-fold higher in young mice compared to aged (**Fig. 1j, Supplementary Fig. 4**). Similarly, Th2 responses (CD3 CD4 IL-4 GATA3) were reduced in aged mice, with a ∼2.47-fold higher frequency in young animals (**Fig. 1k, Supplementary Fig. 5**).

Together, these results confirmed that aging compromises both antibody production and T-cell activation following mRNA LNP vaccination. These results motivate a deeper analysis of where in the vaccination process this deficit arises.

### Early immune cell recruitment, transfection, and antigen presentation remain largely intact in aged mice

To determine whether aging alters the early innate response to mRNA LNP vaccination, we assessed immune cell recruitment 24 h post i.m. injection of 10 μg mOVA SM-102 LNP. Muscle tissue from the injection site was harvested and analysed by flow cytometry. We first quantified the total number of infiltrating immune cells (CD45) and found no significant difference between young and aged mice (**Fig. 2a**; *p =* 0.08). Next, we profiled key innate immune subsets. There were no significant differences between age groups in the abundance of dendritic cells (CD45 CD11b CD19 Ly6G CD11c MHCII cells), macrophage-like cells (CD45 CD11b CD19 Ly6G CD11c cells), neutrophil-like cells (CD45 CD11b CD19 Ly6G cells), or natural killer (NK) cells (CD45 CD11b CD19 NKp46 cells) in the injected muscle (**Fig. 2b–e. Supplementary Fig. 6**). Similarly, adaptive immune cell populations, including B cells and CD8 T cells, were present at comparable levels between young and aged mice, with the exception of CD4 T cells (CD45 CD4), which were modestly reduced in aged mice (*p =* 0.04; **Fig. 2f–h, Supplementary Fig. 7**). These data suggest that immune cell recruitment to the injection site is largely preserved with age at this early time point.

**Figure 2.**
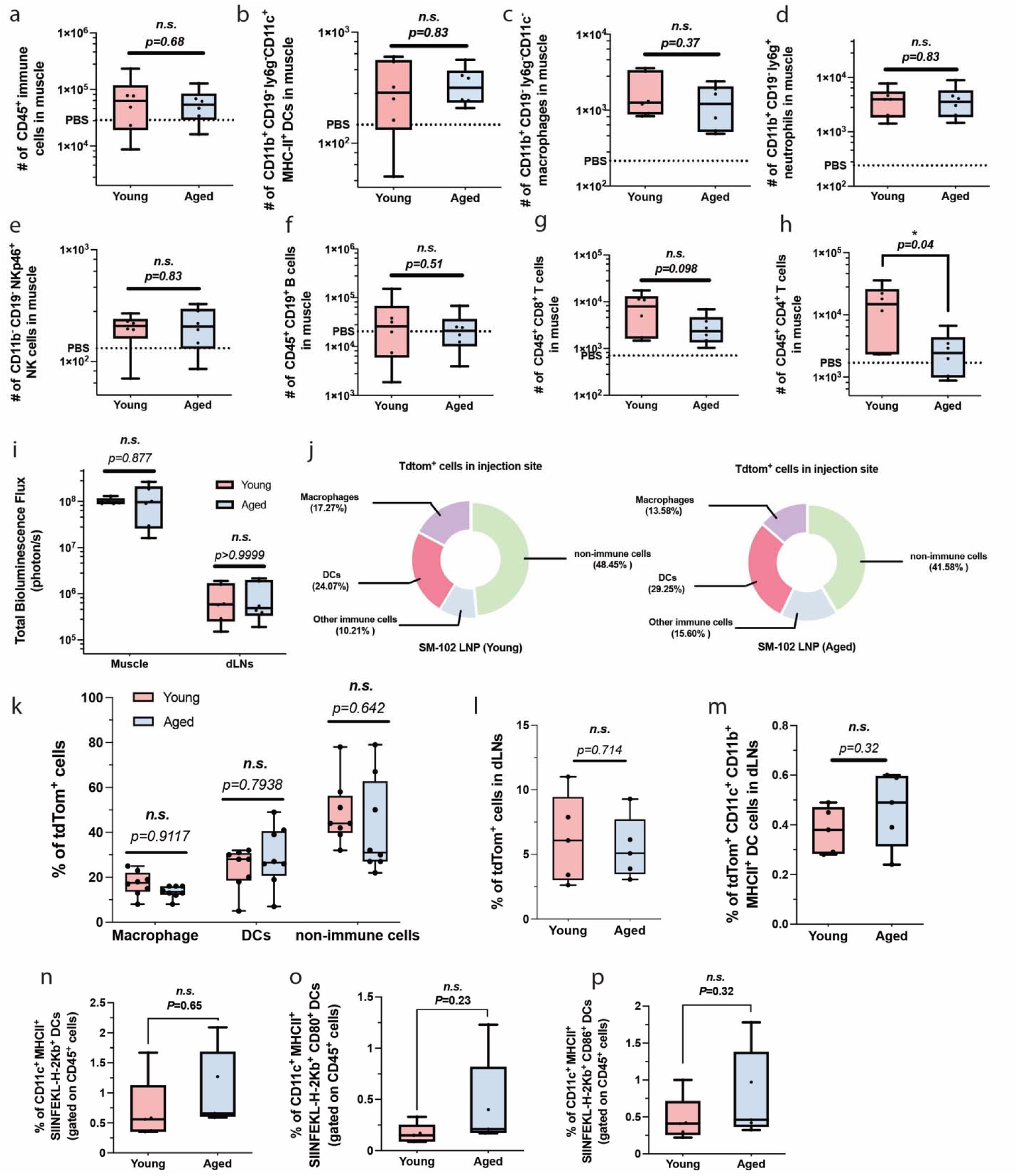
Early immune cell recruitment, transfection, and antigen presentation in the local injection site of young and aged mice following SM-102 mRNA-LNP vaccination. **a**–**h,** Flow cytometry analysis of immune cell infiltration into the injection site (muscle) at 24 h after i.m. administration of 10 μg SM-102 mOVA LNP in young and aged C57BL/6 mice. Muscle tissue (∼1 g) was enzymatically digested, and immune cell subsets were quantified, including total CD45⁺ immune cells (**a**), dendritic cells (CD45⁺CD11b⁺CD19⁻Ly6G⁻CD11c⁺MHCII⁺; **b**), macrophage-like cells (CD45⁺CD11b⁺CD19⁻Ly6G⁻CD11c⁻; **c**), neutrophil-like cells (CD45⁺CD11b⁺CD19⁻Ly6G⁺; **d**), NK cells (CD45⁺CD11b⁻CD19⁻NKp46⁺; **e**), B cells (CD45⁺CD19⁺; **f**), CD8⁺ T cells (CD45⁺CD8⁺; **g**), and CD4⁺ T cells (CD45⁺CD4⁺; **h**). **i**, IVIS of luciferase expression at the injection site and dLNs at 24 h post-injection of mLuc-SM-102 LNPs (10LJμg) in young and aged mice. **j**–**k**, Ai9 Cre-reporter mice were i.m. injected with 10LJμg of mCre-loaded SM-102 LNPs, and transfected cell populations in the muscle were analysed on day 5. Quantification includes tdTomato⁺ non-immune cells (CD45⁻), DC-like populations (CD45⁺ CD11c⁺ CD11b⁺ MHCII⁺ cells), and macrophage-like populations (CD45⁺ CD11c⁻ CD11b⁺ cells). **l**–**m**, Transfection efficiency in dLNs on day 5 post-injection in Ai9 mice, including total tdTomato⁺ cells (**l**) and transfected DC-like cells (tdTomato⁺ CD11c⁺ CD11b⁺ MHCII⁺ cells; **m**). **n**–**p**, Antigen presentation in dLNs was assessed by flow cytometry in young and aged C57BL/6 mice on day 5 post-injection of 10LJμg mOVA-loaded SM-102 LNPs. Quantification includes total antigen-presenting DCs (CD45⁺ CD11c⁺ MHCII⁺ SIINFEKL-H2KLJ⁺ cells; **n**), and expression of co-stimulatory molecules CD80 (**o**) and CD86 (**p**) on antigen-presenting DCs. Data represent mean ± s.e.m. (n = 6 mice per group for **a**–**i**; n = 8 mice per group for **j**–**k**; n = 5 mice per group for **l**–**p**). Data were analysed using an unpaired two-tailed t-test (**a**–**h**, **l**–**p**) or one-way ANOVA with Tukey’s multiple comparisons test (**i**, **k**). \**p* < 0.05; n.s., not significant.

To evaluate whether age affects LNP-mediated transgene expression, we measured luciferase activity in the injection site and dLNs at 24 h post-vaccination using *in vivo* bioluminescence imaging. As shown in **Fig. 2i**, luciferase signal intensity was comparable between young and aged mice in both muscle (*p* = 0.877) and dLNs (*p* > 0.9999), indicating that initial mRNA delivery and translation were unaffected by aging.

Given previous reports that the identity of transfected cells influences immune outcomes, we employed the Ai9 Cre-reporter mouse model to characterize the cellular profile of transfected cells five days post-vaccination. Flow cytometric analysis revealed a similar distribution of tdTomato cells between young and aged mice (**Fig. 2j–k, Supplementary Fig. 8**). In both groups, about 40–50% of transfected cells were non-immune (CD45), while among CD45 transfected cells, DC-like cells represented about 20–30%. No significant differences were observed in the transfection rates of DC-like or macrophage-like cells. We further analysed tdTomato cells in the dLNs and again found no significant difference between young and aged mice in the frequency of transfected cells (*p =* 0.714; **Fig. 2l, Supplementary Fig. 8**) or transfected DCs (tdTomato CD11c CD11b MHCII cells; *p =* 0.32; **Fig. 2m, Supplementary Fig. 8**). Finally, we assessed antigen presentation five days post-vaccination by quantifying the frequency of CD11c MHCII SIINFEKL-H2Kb DCs in the dLNs. There were no significant differences between young and aged mice in the frequency of antigen-presenting DCs (**Fig. 2n, Supplementary Fig. 9**), nor in the proportion expressing co-stimulatory molecules CD80 and CD86 (**Fig. 2o–p, Supplementary Fig. 9**).

These results show that early local immune cell recruitment, transgene expression, and antigen presentation at the injection site and dLNs proceed efficiently in aged hosts, suggesting that age-associated defects in vaccine efficacy arise downstream of these initial events.

### Aging alters immune cell gene expression at the injection site and reduces systemic mRNA-LNP transfection efficiency

Although cellular responses at the injection site appeared comparable between young and aged mice, transcriptomic profiling revealed molecular differences in immune activation. CD45 immune cells were isolated from the injection site 24 h after i.m. administration of SM-102 mOVA LNP (10 μg per mouse) using a Percoll density gradient, and RNA was extracted and profiled using a Nanostring immunology panel. In young mice, 99 genes were significantly altered compared to PBS controls, with 74 upregulated and 25 downregulated (**Fig. 3a**). In aged mice, only 53 genes were differentially expressed (26 upregulated, 27 downregulated) (**Fig. 3b**). Notably, direct comparison between young and aged mice revealed 83 differentially expressed genes, of which 79 were upregulated in young animals, suggesting a marked attenuation of immune activation in aged mice (**Fig. 3c**).

**Figure 3.**
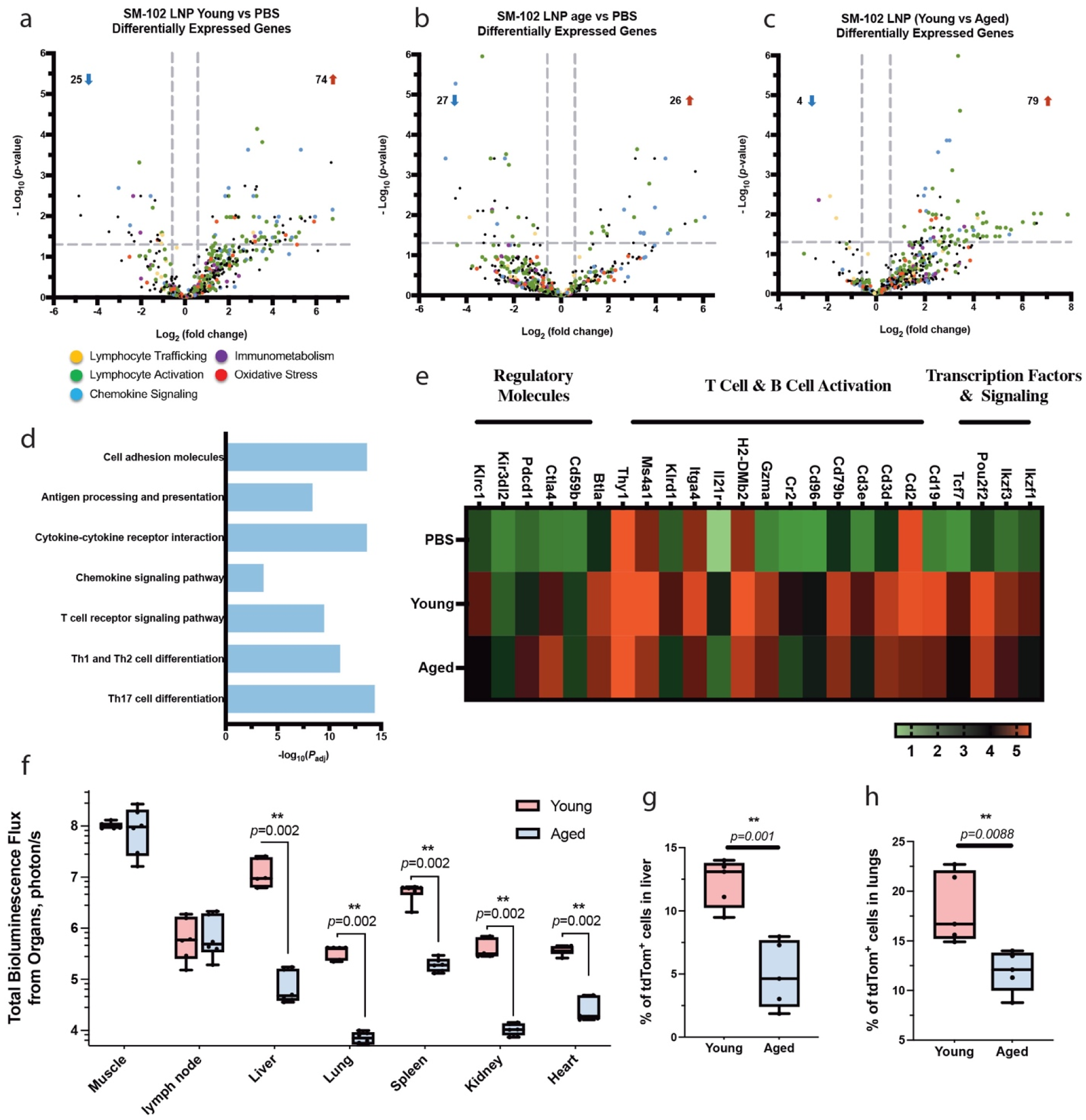
Aged mice exhibit impaired immune gene activation and systemic transgene expression following SM-102 mRNA-LNP vaccination. **a**–**c,** Immune cells were isolated from the injection site 24 h after i.m. administration of SM-102 mOVA LNPs (10 μg per mouse) in young and aged C57BL/6 mice. RNA was extracted and analysed using a Nanostring immunology panel. Volcano plots show differentially expressed genes in young mice versus PBS (**a**), aged mice versus PBS (**b**), and direct comparison between young and aged mice (**c**) (two-sided unpaired limma-moderated t-test; absolute fold change ≥ 1.5, *p* < 0.05). **d,** KEGG pathway enrichment analysis (via DAVID) comparing gene expression profiles of immune cells from young and aged mice revealed downregulation of key immune activation pathways in aged animals. **e,** Heatmap showing expression levels of selected genes involved in antigen presentation, T and B cell activation, regulatory molecules, and transcription factors in immune cells from the injection site of young and aged mice. **f,** IVIS of luciferase expression 24 h post-vaccination with mLuc-loaded SM-102 LNPs demonstrates significantly reduced systemic transgene expression in aged mice across major organs, including the liver, lungs, and spleen. **g**–**h**, Ai9 Cre-reporter mice were used to evaluate transfection efficiency in peripheral organs 5 days post i.m. injection of mCre-loaded SM-102 LNPs (10 μg per mouse). Quantification of tdTomato⁺ cells in the liver (**g**) and lungs (**h**) confirms impaired systemic cellular transfection in aged mice. Data represent mean ± s.e.m. (n = 3 mice per group for **a**–**e**; n = 6 mice per group for **f**–**h**). Data were analysed using two-way ANOVA and Tukey’s multiple comparisons test (**f**) or unpaired two-tailed t-tests (**g**–**h**). \*\**p* < 0.01.

To interpret these transcriptional differences in the context of immune function, we conducted Kyoto Encyclopedia of Genes and Genomes (KEGG) pathway analysis using the DAVID database. Aged mice showed lower expression of genes involved in antigen processing and presentation, cytokine–cytokine receptor interactions, chemokine signalling, T cell receptor signalling, and Th1/Th2 cell differentiation pathways (**Fig. 3d, e**). These results suggest that, despite comparable immune cell infiltration and transfection at the injection site, immune cells from aged mice exhibit functional deficiencies in immune cell transcriptional programs.

We next examined systemic transgene expression, given prior evidence that mRNA LNP vaccines can traffic to distal organs such as the liver, lungs, and spleen, driving tissue-specific immune responses, including tissue-resident memory T cells (Trm).^32,33^ At 24 h post-vaccination, *in vivo* bioluminescence imaging revealed stark differences in systemic luciferase expression between young and aged mice (**Fig. 3f**). While local signal intensities at the injection site and dLNs were similar, aged mice showed dramatically reduced transgene expression in peripheral organs: a ∼162-fold reduction in the liver, ∼49-fold in the lungs, and ∼27-fold in the spleen compared to young animals. To confirm these findings at the cellular level, we used Ai9 Cre-reporter mice to assess transfection in individual organs at day 5 post-injection. In the liver, only 4.9% of cells were tdTomato in aged mice, compared to 12.0% in young mice (**Fig. 3g**). A similar reduction was observed in the lungs, where transfection efficiency dropped from ∼18.3% in young mice to ∼11.9% in aged mice (**Fig. 3h**).

To determine whether these differences in transgene expression were due to altered biodistribution, we administered Cy5-labeled mRNA-loaded SM-102 LNPs and assessed organ fluorescence at 24 h. Cy5 signal was comparable across young and aged mice in all major organs, including the liver and lungs, indicating that LNP transport and distribution were not impaired by age (**Supplementary Fig. 10**). This suggests that LNP trafficking was intact and that the reduced overall transgene expression stems from intrinsic age-related differences in cellular uptake, endosomal escape, or translational capacity.

Together, these results show that aging dampens transcriptional activation of immune cells at the injection site and markedly reduces systemic mRNA translation in key immune-relevant organs, while leaving early local processes intact. These molecular and functional deficits point to systemic translation as a previously underappreciated point of vulnerability in mRNA-LNP vaccine performance.

### Restoring systemic mRNA expression rescues antigen-specific immune responses in aged mice

Given our observation that systemic transgene expression is markedly reduced in aged mice following *i.m.* administration of mRNA-LNP vaccines, we next test whether this deficiency contributes directly to the attenuated immune responses observed with age. We used an intravenous (*i.v.*) ‘boost’ strategy designed to restore systemic mRNA expression in aged animals. Specifically, aged mice received a standard i.m. dose of SM-102 mOVA LNP (10 μg mRNA) followed immediately by an additional *i.v.* dose of 3 μg mRNA LNPs. *In vivo* imaging at 24 h post-administration revealed that this combined dosing strategy successfully restored transgene expression in peripheral organs of aged mice to levels comparable to those in young mice receiving *i.m.* vaccination alone. Luciferase signal intensity in the liver (*p =* 0.9897), lungs (*p* > 0.9999), and spleen (*p* > 0.999) in aged animals was effectively brought to the same level as that in young mice (**Fig. 4a**).

**Figure 4.**
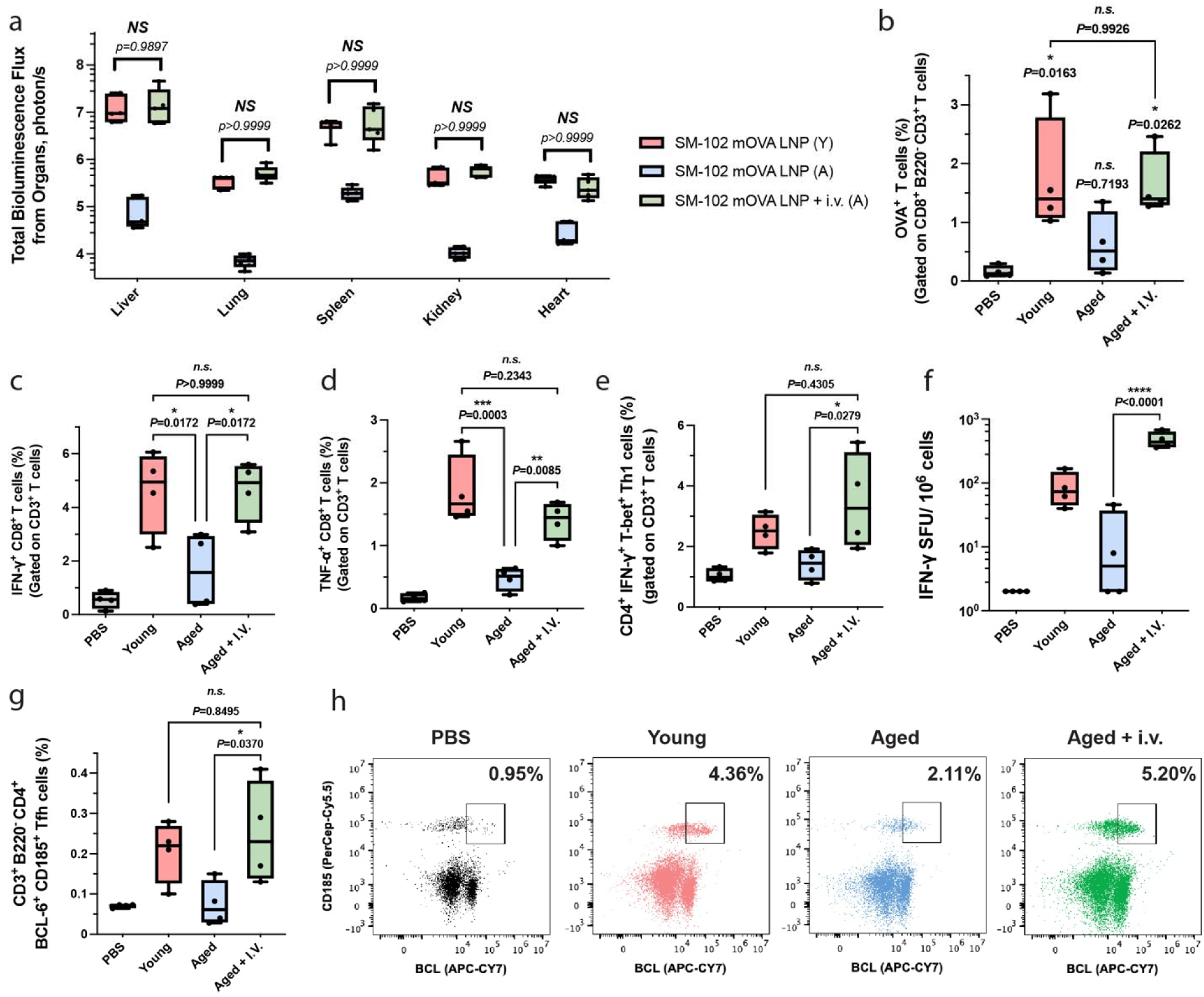
Intravenous mRNA-LNP dosing restores systemic transgene expression and antigen-specific immune responses in aged mice. **a,** To overcome age-associated reductions in systemic mRNA expression, aged mice received a standard i.m. dose of SM-102 mOVA LNPs (10LJμg) followed immediately by an i.v. dose of 3LJμg LNPs. IVIS was performed 24 h post-injection to assess transgene expression in peripheral organs. **b–h**, Mice were vaccinated intramuscularly with 10LJμg of mOVA-loaded SM-102 LNPs on days 0, 7, and 14, and sacrificed on day 30 for immune profiling. **b**, Frequency of OVA-specific CD8⁺ T cells in spleens. **c–e**, Splenocytes were restimulated *ex vivo* with OVA (100LJμgLJmL⁻¹) and SIINFEKL peptide (2LJμgLJmL⁻¹) for 12 h, and cytokine-producing CD8⁺ T cells were analysed by intracellular cytokine staining: IFN-γ⁺ (**c**), TNF-α⁺ (**d**), and CD3⁺CD4⁺Tbet⁺IFN-γ⁺ Th1 cells (**e**). **f**, IFN-γ FluoroSpot assay of restimulated splenocytes. **g**–**h**, Frequency of CD3⁺B220⁻CD4⁺BCL6⁺CD185⁺ follicular helper T (Tfh) cells in the spleen. Data represent mean ± s.e.m. (n = 5 mice per group for **a**, n = 4 mice per group for **b**–**h**). Data were analysed using two-way ANOVA and Tukey’s multiple comparisons test (**a**) and unpaired t-tests (**b**–**h**). Exact *p*-values are shown; \**p <* 0.05, \*\**p <* 0.01, \*\*\*\**p <* 0.0001, *n.s.,* not significant.

Using a prime-boost-boost schedule on days 0, 7, and 14, we evaluated immune responses on day 30. Notably, aged mice that received an additional *i.v.* dose exhibited a significant increase in OVA-specific CD8 T cells in the spleen, ∼2.59-fold higher than aged mice that received i.m. vaccination alone, and statistically indistinguishable from the young that received only a standard i.m. dose cohort (*p =* 0.9926; **Fig. 4b, Supplementary Fig. 1**). Cytokine-producing cytotoxic T cells in the spleen were similarly restored. The percentage of IFN-γ CD8 T cells was ∼2.84-fold higher in the i.m. + i.v. group in aged mice, compared to i.m.-only aged mice, matching levels in young mice received a standard i.m. dose alone (*p* > 0.9999; **Fig. 4c, Supplementary Fig. 2**). TNF-α CD8 T cells also increased ∼2.95-fold with the *i.m.* + *i.v.* dosing schedule in aged mice approached the young cohort baseline (*p =* 0.2343; **Fig. 4d**). We further assessed antigen-specific CD4 T cell responses in the spleen. The proportion of Th1-polarized cells (CD3 CD4 IFN-γ Tbet cells) was elevated ∼2.48-fold in aged mice receiving the i.v. boost, again approximating levels observed in young mice (*p =* 0.4305; **Fig. 4e, Supplementary Fig. 4**). Consistent with these findings, FluoroSpot analysis revealed a dramatic ∼33-fold increase in IFN-γ–secreting splenocytes compared to i.m.-only aged mice (**Fig. 4f**). Finally, we examined antigen-specific follicular helper T cell (Tfh) responses in the spleen. Aged mice that received the *i.v.* supplement dose showed a ∼3.33-fold increase in the frequency of antigen-specific Tfh cells (CD3 B220 CD4 BCL6 CD185 cells), again reaching levels comparable to the i.m.-only young group (*p =* 0.8495; **Fig. 4g, h, Supplementary Fig. 11**).

Together, these data demonstrate that restoring systemic transgene expression by supplementing i.m. vaccination with a low-dose *i.v.* mRNA LNP injection is sufficient to rescue both CD8+ and CD4+ T cell responses in aged mice. These findings highlight the critical role of systemic transfection in driving robust immune responses and suggest that impaired LNP-mediated mRNA delivery to peripheral organs is a key bottleneck in aged hosts.

### Restored systemic transgene expression reinstates antitumor efficacy and sustains antigen-specific T cell responses in aged mice

We next examined whether enhanced systemic transgene expression could translate into rescued immune responses and therapeutic efficacy in aged mice. In a therapeutic B16-OVA melanoma model, young and aged mice received OVA mRNA (mOVA)-loaded SM-10 LNPs (10 μg mOVA per dose, *i.m.*) on days 4, 11, and 18 following subcutaneous B16-OVA tumour inoculation (**Fig. 5a**). As expected, aged mice receiving the *i.m.* vaccination alone exhibited accelerated tumour progression and shortened median survival relative to young controls (*p =* 0.0072, **Fig. 5b, c**). Strikingly, co-administration of a low-dose (3 μg mOVA) *i.v.* LNPs with each i.m. vaccination fully restored systemic transgene levels and rescued tumour control in aged mice. Tumour growth kinetics and survival in *the i.m.* + *i.v.* vaccination in aged group were comparable to those in young counterparts (tumor volume, *p =* 0.6389; survival, *p =* 0.7921).

**Figure 5.**
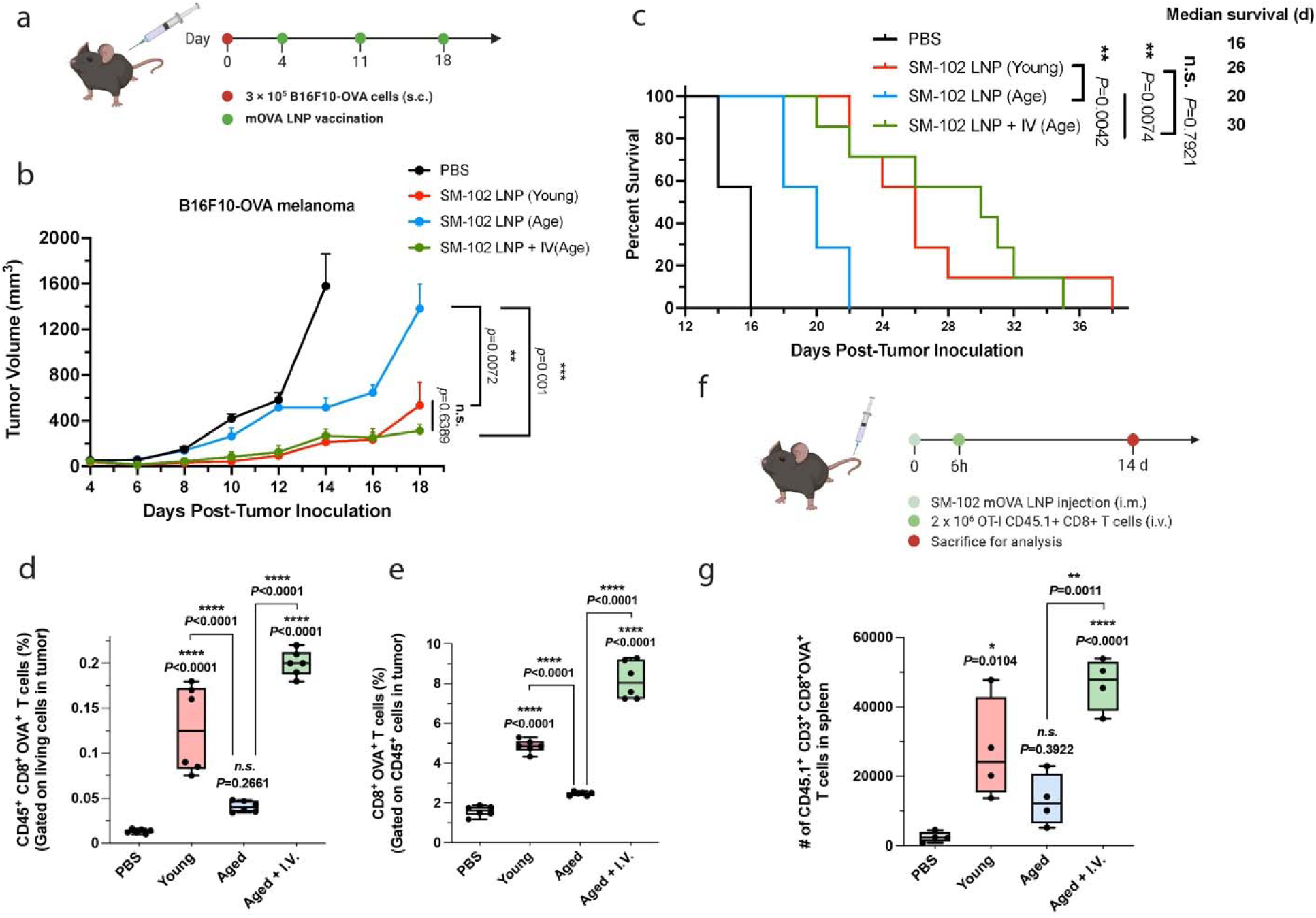
Restored systemic transgene expression reinstates antitumour efficacy and sustains antigen-specific T cell responses in aged mice. **a,** Schematic of the therapeutic vaccination model. Mice were subcutaneously inoculated with B16-OVA tumour cells on day 0, followed by mRNA SM-102 LNP vaccination (10 μg mOVA, i.m.) on days 4, 11, and 18. **b, c,** Tumour growth (**b**) and survival curves (**c**) following vaccination. **d, e,** Analysis of tumour-infiltrating lymphocytes on day 14 post-tumour inoculation. Tumours were dissociated into single-cell suspensions and analysed by flow cytometry. Shown are the frequency of CD45⁺CD8⁺OVA⁺ T cells among total live cells (**d**) and the frequency of CD8⁺OVA⁺ T cells among CD45⁺ live cells (**e**). **f,** Schematic of adoptive transfer experiment. Mice received mOVA SM-102 LNPs (10 μg mOVA, i.m. with or without 3 μg mOVA i.v.). Six hours later, 2 × 10LJ CD45.1⁺ OT-I CD8⁺ T cells were injected intravenously. Splenic OT-I T cell levels were assessed two weeks post-transfer. **g**, Number of CD45.1⁺CD3⁺CD8⁺OVA⁺ T cells in spleens collected 14 days post-transfer. Data represent mean ± s.e.m. (n = 7 mice per group for **b–c**, n = 6 mice per group for **d–e**, n = 4 mice per group for **g**). Data were analysed using two-way ANOVA and Tukey’s multiple comparisons test (**b, d, e, g**). Survival curves (**c**) were compared using the log-rank Mantel–Cox test with Holm–Šídák correction for multiple comparisons. \**p* < 0.05, \*\**p* < 0.01, \*\*\**p* < 0.001, \*\*\*\**p* < 0.0001; n.s., not significant. Panels **a** and **f** were created in BioRender.

To further investigate the immunological basis of this restored tumour control, we examined the tumour microenvironment on day 14 post-tumour inoculation. Tumours were collected, processed into single-cell suspensions, and analysed via flow cytometry to assess antigen-specific T cell infiltration. Aged mice received only i.m. vaccination exhibited lower levels of OVA-specific CD8 T cells than young mice. Specifically, the frequency of CD45 CD8 OVA T cells among total live cells was ∼3.11-fold lower, and the frequency of CD8 OVA T cells among CD45 live cells was ∼1.96-fold lower (**Fig. 5d, e**). In contrast, aged mice that received the i.m. + i.v. vaccination displayed a substantial enhancement in tumour-infiltrating OVA-specific CD8 T cells compared to the young cohort. The percentage of CD45 CD8 OVA T cells among live cells was 1.57-fold higher, and the percentage of CD8 OVA T cells among CD45 live cells was 1.68-fold higher than the *i.m.* only group, reaching levels comparable to or exceeding those in the young cohort.

We next explored whether sustained systemic antigen expression contributes to the restored antigen-specific T cell responses. Previous studies suggest that antigen re-encounter, especially in peripheral tissues, plays an important role in memory and effector T cell persistence. We designed an adoptive transfer experiment in which mice received either *i.m.* alone or *i.m. + i.v.* mOVA LNP vaccination, followed by *i.v.* injection of 2 × 10 CD45.1 OT-I CD8 T cells 6 h later. Two weeks post-transfer, splenic OT-I T cell levels were analysed to assess persistence and recall (**Fig. 5f**). In PBS control group, OT-I cells were nearly undetectable at the two-week timepoint, highlighting the necessity of antigen exposure for T cell maintenance (**Fig. 5g, Supplementary Fig. 12**). Among vaccinated groups, aged mice with *i.m.* vaccination showed a ∼2.09-fold reduction in CD45.1 CD3 CD8 OVA T cells compared to young mice. Aged mice received the *i.m.* + *i.v.* vaccination exhibited a ∼2-fold higher splenic OT-I T cells than young mice with *i.m.* vaccination (**Fig. 5e, Supplementary Fig. 12**). These findings suggest that systemic antigen expression plays a key role not only in priming but also in sustaining antigen-specific CD8 T cells.

Together, these findings demonstrate that rescuing systemic transgene expression restores therapeutic efficacy and maintains antigen-specific T cell responses in aged hosts. Persistent antigen expression in peripheral organs appears essential for maintaining antigen-specific T cells, providing a mechanistic explanation for how systemic mRNA translation governs age-dependent vaccine efficacy.

### LNP composition governs systemic transgene expression and enables age-independent immune response and vaccine efficacy

Building on our findings that restored systemic transgene expression restores antitumour efficacy in aged mice, we next investigated whether formulation tuning could intrinsically overcome the constraint in systemic translation and immune response observed in aged hosts. Given that LNP composition critically influences nanoparticle trafficking, biodistribution, and transfection efficiency, we screened five previously characterized LNP formulations (A–E; **Supplementary Table 1**) from our prior studies^18,19,33^ for their ability to mediate systemic transgene expression following *i.m.* injection in young mice. Among these, Formulation B (LNP B) exhibited a 2.68-fold higher luciferase expression in the liver at 24 h post-injection compared to the SM-102 LNPs (**Fig. 6a**).

**Figure 6.**
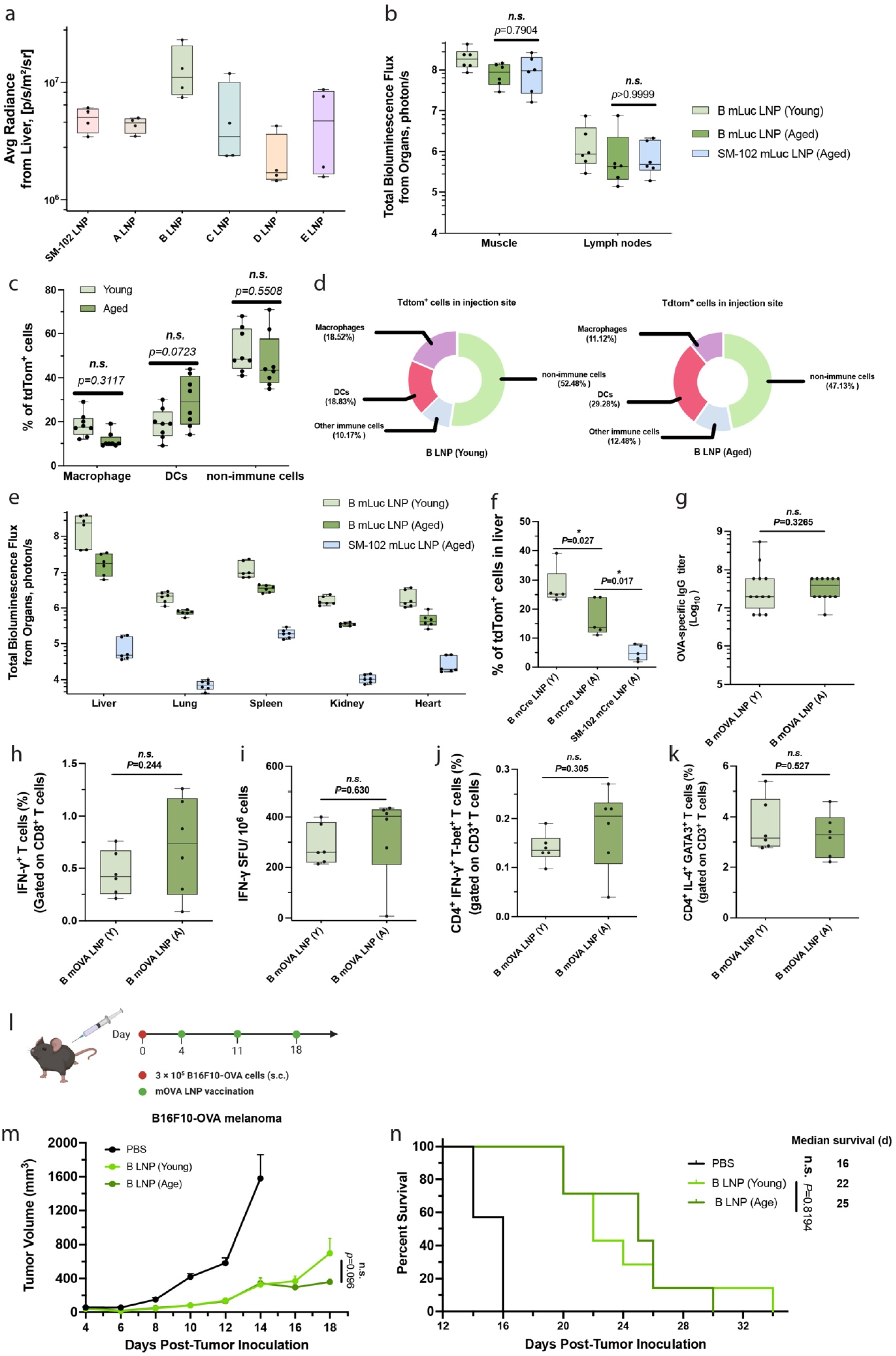
LNP composition influences systemic transgene expression and immune responses in aged mice. **a,** Screening of previously characterized mLuc-loaded LNP formulations (A–E, 10LJμg per mouse; **Supplementary Table 1**) in young mice. IVIS imaging was used to quantify luciferase expression in the liver at 24LJh after i.m. injection. **b**, Luciferase expression at the injection site and dLNs in young and aged mice 24LJh post-i.m. delivery of LNP B. **c**–**d**, Analysis of transfected cell populations at the injection site five days post-vaccination using Ai9 reporter mice. Transfected (tdTomato⁺) cells were quantified among non-immune cells (CD45⁻), DC-like cells (CD45⁺CD11c⁺CD11b⁺MHCII⁺), and macrophage-like cells (CD45⁺CD11c⁻CD11b⁺). **e**, Organ-specific systemic transfection following i.m. injection of LNP B in young and aged mice and SM-102 LNPs in aged mice. Luciferase expression was assessed in the liver, lungs, spleen, kidneys, and heart at 24LJh post-injection. **f**, Quantification of tdTomato⁺ cells in the liver of Ai9 mice five days post-i.m. injection with mCre-loaded LNP B. **g–k**, Young and aged C57BL/6 mice were vaccinated i.m. with 10LJμg mOVA-loaded LNP B on days 0, 7, and 14, and analysed on day 30. Serum OVA-specific IgG titres (**g**), IFN-γ⁺ CD8⁺ T cells (**h**), IFN-γ FluoroSpot responses (**i**), CD3⁺CD4⁺IFN-γ⁺Tbet⁺ Th1 cells (**j**), and CD3⁺CD4⁺IL-4⁺GATA3⁺ Th2 cells (**k**) were measured from restimulated splenocytes. **l**, Schematic of the therapeutic vaccination model. Mice were subcutaneously inoculated with B16-OVA tumour cells on day 0, and vaccinated with 10LJμg mOVA-loaded LNP B (i.m.) on days 4, 11, and 18. **m–n**, Tumour growth (**m**) and survival curves (**n**) in young and aged mice following vaccination with LNP B. Data represent mean ± s.e.m. (n = 4 mice per group for **a**, n = 6 mice per group for **b, e, h–k**, n = 8 mice per group for **c–d**, n = 5 mice per group for **f**, n = 12 mice per group for **g**, n = 7 mice per group for **m–n**). Data were analysed using two-way ANOVA and Tukey’s multiple comparisons tests (**b, c, f, m**), unpaired t-tests (**g–k**). Survival (**n**) was analysed using the log-rank Mantel–Cox test with Holm–Šídák correction for multiple comparisons. \**p* < 0.05; n.s., not significant. Panel **l** was created in BioRender.

This LNP B induced comparable local transfection in muscle at the injection site (slightly lower in aged mice; *p =* 0.0272) and in dLNs (*p* > 0.9999) between the young and aged groups (**Fig. 6b**), and similar transgene expression levels to the SM-102 LNPs in aged mice at both the injection site (*p =* 0.7904) and dLNs (*p* > 0.9999). Analysis of transfected cell profiles at the injection site using Ai9 reporter mice on day 5 post-vaccination revealed that LNP B produced a similar cellular distribution on young and age mice: approximately 50% of transfected cells were non-immune (CD45) (*p =* 0.5509), and among immune cells, ∼20% were DCs (*p =* 0.0723) in both young and aged mice (**Fig. 6c, d**).

We then compared local and systemic expression profiles of LNP B in aged and young mice. Age-associated reduction in transgene expression was also observed with LNP B: 11.5-fold in the liver, 2.76-fold in the lungs, and 3.59-fold in the spleen. Nonetheless, these reductions were markedly smaller than those observed with SM-102 LNPs (**Fig. 6e**). Compared to SM-102 LNPs in aged mice, LNP B achieved markedly higher systemic expression: 229-fold in the liver, 105-fold in the lungs, and 18-fold in the spleen. These results were corroborated using Ai9 mice five days post-vaccination with mCre-loaded LNP B. In the liver, 27.6% of cells were tdTomato in young mice, while 17.2% were transfected in aged mice, substantially higher than the 4.95% observed in aged mice that received SM-102 LNPs (**Fig. 6f**). To assess biodistribution, we delivered Cy5-labeled mRNA via LNP B and imaged major organs at 24 h post-injection. We observed no significant differences between young and aged mice, consistent with biodistribution results from SM-102 LNPs, further confirming that the overall trafficking pattern of LNPs remains intact with age and that the age-associated reduction in systemic transfection arises from compromised intracellular processing rather than altered delivery routes (**Supplementary Fig. 13**).

To evaluate whether the enhanced systemic expression translated into functional benefits, we compared immune responses induced by LNP B in young and aged mice using the same vaccination schedule as showed in **Fig. 1a**. No significant differences were observed between young and age groups in OVA-specific IgG titres (*p =* 0.3265), IFN-γ CD8 T cells (*p =* 0.244), IFN-γ FluoroSpot responses (*p =* 0.63), Th1-polarized CD4 T cells (*p =* 0.305), or Th2-polarized CD4 T cells (*p =* 0.527) (**Fig. 6g–k, Supplementary Fig. 14**). In the therapeutic B16-OVA melanoma model where mice were inoculated with tumour cells on day 0 and treated with three doses of mOVA LNP B on days 4, 11, and 18 (**Fig. 6l**), tumour growth and survival were indistinguishable between young and aged mice (*p =* 0.096 for tumour growth, *p =* 0.8194 for survival; **Fig. 6m–n, Supplementary Fig. 15**).

Together, these findings demonstrate that tuning LNP composition can minimize age-associated disparities in systemic mRNA translation and immune activation. The mRNA LNP formulations can be optimized to achieve robust systemic expression across age groups without altering the administration route, thereby overcoming translational senescence and maintaining vaccine efficacy in aged hosts. This study provides an effective strategy for the rational design of age-adaptive mRNA LNP formulations capable of delivering consistent immune and therapeutic outcomes across age groups.

## DISCUSSION

Aging is widely recognized to impair immune responses to vaccination, but the underlying mechanisms remain incompletely defined. Clinical studies of COVID-19 mRNA vaccines illustrate this challenge: in individuals over 80 years of age, approximately half fail to generate robust neutralizing antibody responses to the BNT162b2 vaccine, accompanied by lower frequencies of spike-specific memory B cells and diminished IL-2–producing CD4 T cells.^26^ These data underscore both quantitative and qualitative immune impairments with age. However, human analyses have been largely confined to peripheral blood measurements, providing limited insight into antigen presentation, lymphoid tissue responses, and vaccine biodistribution—all critical determinants of mRNA vaccine performance.^25–27,29^

In this study, we systematically examined how aging alters each step of the mRNA LNP vaccination cascade. We found that aged mice exhibit markedly reduced humoral and cellular immune responses, including IgG titres, cytotoxic CD8 T cell responses, and Th1/Th2 activation, compared to young counterparts, consistent with clinical observations.

Yet, the magnitude and composition of early local events, including immune cell recruitment at the injection site, transfection in local tissue and dLNs, and antigen presentation and DC maturation, remained largely intact. These results indicate that classical features of immunosenescence, such as defective antigen-presenting cell activation, do not fully explain the age-related decline in mRNA vaccine efficacy.

Instead, our data reveal that systemic mRNA translation in distal organs, including the liver, lungs, and spleen, was markedly reduced in aged mice. This defect occurs despite equivalent biodistribution of LNPs and cellular uptake, suggesting that aging perturbs intracellular processing or translation rather than delivery per se. Restoration of systemic transgene expression in aged mice either through i.v. co-administration or through LNP formulation tuning, fully rescues both T cell immunity and tumour control in aged mice. These findings position systemic mRNA translation as a previously underappreciated determinant of LNP vaccine efficacy.

At the molecular level, transcriptomic profiling of immune cells from the aged mice revealed attenuated expression of genes associated with antigen processing and presentation, T cell activation and differentiation, and chemokine signalling. These transcriptional changes likely contribute to the reduced vaccine efficacy in aged animals.

These cellular and molecular signatures are consistent with emerging evidence that aging alters global protein synthesis and translation efficiency, a phenomenon sometimes referred to as translational senescence, characterized by reduced ribosome biogenesis and dysregulation of mRNA translation. Although further studies will be required to directly test these mechanisms, such age-associated changes could limit the efficiency with which LNP-delivered mRNA vaccine is processed and translated into antigenic proteins in local tissue and systemic organs following *i.m.* immunization.

Beyond providing mechanistic insight, these findings have important translational implications. Our results demonstrate that age-associated declines in mRNA vaccine efficacy are not inevitable but can be overcome by engineering LNP formulations that sustain robust systemic mRNA translation. The LNP B formulation identified here exemplifies this concept, achieving age-independent immune activation and therapeutic efficacy without altering the route of administration. Together, these data outline a rational path toward age-adaptive mRNA vaccines, formulations intentionally tuned to the metabolic and translational landscape of the aging host, with potential relevance not only for cancer immunotherapy but also for prophylactic vaccination and mRNA-based treatments for infectious and chronic diseases of aging.

An additional question raised by this work is whether the optimization strategies identified here are specific to the benchmark SM-102 LNP or extend more broadly to other clinically relevant mRNA LNP platforms. Although our mechanistic analyses centered on SM-102 as a well-characterized, commercially deployed formulation, multiple observations support the generalizability of our conclusions. Age-associated loss of systemic transgene expression occurred without changes in biodistribution across two chemically distinct LNPs, indicating that impaired intracellular processing or translation, rather than delivery, is the dominant age-sensitive step. Moreover, LNP B, which differs substantially from SM-102 in both ionizable lipid chemistry and helper lipid composition and was not designed with aging in mind, exhibited enhanced systemic expression and fully eliminated age-dependent disparities in immune response and antitumor efficacy. These results suggest that the critical, tunable determinant is an LNP’s capacity to sustain mRNA translation in aged tissues, rather than reliance on any single proprietary lipid. While we did not systematically evaluate all clinically deployed LNPs, the convergence of findings across distinct chemistries supports systemic translation efficiency as a broadly applicable, formulation-agnostic lever for improving mRNA vaccine performance in older hosts.

In summary, this study establishes systemic mRNA translation as a tunable determinant of age-dependent vaccine efficacy. While early local immune activation is preserved with age for mRNA LNP vaccines, downstream deficits in mRNA translation constrain adaptive immunity. LNP formulation optimization targeting restoration of systemic transgene expression offers a promising strategy to enhance mRNA vaccine performance in the populations who stand to benefit most.

## METHODS

### Ethical statement

All animal experiments were performed in accordance with relevant ethical guidelines and were approved by the Johns Hopkins University Institutional Animal Care and Use Committee (IACUC) under protocol number MO24E165.

The IACUC-approved maximum tumor burden was 2,000 mm³ or a maximum diameter of 20 mm in any dimension. Mice were euthanized when either limit was reached. We confirm that these thresholds were not exceeded in any experiments described in this study.

### Materials

Ionizable lipids including SM-102 (Cat#BP-25499), ALC-0315 (Cat#BP-25498) and DLin-MC3-DMA (Cat# BP-25497) was purchased from Broadpharm. DOPE, DSPC, 18PG, and DMG-PEG-2000 were obtained from Avanti Polar Lipids. Cholesterol was from Sigma-Aldrich. B16-OVA cells were kindly provided by the lab of Prof. Jonathan P. Schneck. All eukaryotic cell lines used in this study were authenticated by short tandem repeat profiling and routinely tested for mycoplasma contamination using the Universal Mycoplasma Detection Kit (ATCC 30-1012K) every two months. No contamination was detected during the study period. All mRNA (Cre mRNA and OVA mRNA) constructs were purchased from TriLink BioTechnologies.

### LNP synthesis and characterization

LNPs were synthesized by directly adding an organic phase containing the lipids to an aqueous phase containing mRNA. To prepare the organic phase, a mixture of ionizable lipids (SM-102, ALC-0315 or DLin-MC3 DMA), cholesterol, DMG-PEG2000, and a helper lipid (DOPE, DSPC, or 18PG) were dissolved in ethanol. The mRNA (Cre mRNA and OVA mRNA) was dissolved in 25 mM magnesium acetate buffer (pH 4.0). For larger scale LNP production, the aqueous and ethanol phases prepared were mixed at a 3:1 ratio in a flash complexation (FNC) device using syringe pumps and purified by dialysis against DI water using a 100-kDa MWCO cassette at 4 °C for 24 h and were stored at 4 °C before injection. The size, polydispersity index, and zeta potentials of LNPs were measured using dynamic light scattering (ZetaPALS, Brookhaven Instruments). Diameters are reported as the intensity average.

### Animals

Male and female C57BL/6 mice (6–8 weeks old for the young group, 10–12 months old for the aged group) were purchased from the Jackson Laboratory and housed under standard conditions in compliance with institutional policies. Male and female Ai9 mice (6–8 weeks for the young group; 10–12 months for the aged group) were bred and aged in the Johns Hopkins Animal Facilities. Mice were randomly assigned to experimental groups. The mice were supplied with free access to pelleted feed and water. The pelleted feed generally contained 5% fibre, 20% protein, and 5–10% fat. The mice usually ate 4–5 g of pelleted feed (120 g per kg body weight) and drank 3–5 mL of water (150 mL per kg body weight) per day. The temperature of the mouse rooms was maintained at 18–26°C (64–79°F) at 30–70% relative humidity with a minimum of 10 room air changes per hour. Standard shoebox cages with corncob as bedding were used to house the mice. The LNPs were given through *i.m.* injection at a predetermined dose per mouse.

### Tissue processing and cell isolation

For isolation of cells from the liver, lungs, and spleen in the Ai9 mouse experiments, the harvested tissues were placed onto 40-μm cell strainers and digested mechanically with the back of a 3-mL syringe plunger in PBS. The cells were pelleted at 300 ×g for 5 min at 4 °C, followed by resuspension in the ACK lysis buffer and incubation at room temperature for 7 min to lyse red blood cells. Cells were then pelleted by centrifugation at 300 ×g for 5 min at 4 °C, washed with PBS, and pelleted twice before staining for flow cytometry. All steps were performed protected from light.

For isolation of lymphocytes from the spleen, harvested spleens were placed onto 40-μm cell strainers. The spleens were mechanically digested through the cell strainers with the back of a 3-mL syringe plunger in a lymphocyte separation medium (PromoCell). The resulting single-cell cell suspensions were transferred to sterile 15-mL tubes, and 1 mL of RPMI-1640 media was slowly added to each tube to form visible layers. The tubes were then centrifuged at 800 ×g for 25 min at 4 °C without brake. Following centrifugation, the middle layer containing lymphocytes was carefully collected and transferred to fresh 15-mL tubes, and 5 mL of RPMI-1640 media was added and mixed thoroughly. The cells were pelleted by centrifugation at 300 ×g for 5 min at 4 °C. The pellet was then resuspended in 3 mL of RPMI-1640 media and centrifuged again at 300 ×g for 5 min at 4 °C. Isolated lymphocytes were suspended in RPMI-1640 media for subsequent analyses.

### Antibodies and staining for flow cytometry

Antibody panels are provided in **Supplementary Table 2**. LIVE/DEAD fixable dead cell stain kits were used to determine the viability of cells. eBioscience Foxp3/Transcription Factor Staining buffer set (ThermoFisher, 00-5523-00) was used for intracellular staining. Isolated cells from the tissues, as described in the previous section, were resuspended and pelleted in 100 µL of antibodies diluted in flow cytometry staining buffer obtained from eBioscience™. The cells were then incubated on ice in the dark for 1 h. After the incubation period, the stained cells were washed twice with PBS and subsequently resuspended in 200 µL of eBioscience™ flow cytometry staining buffer for flow cytometry analysis. Flow data was acquired using an Attune NXT flow cytometer and analysed with FlowJo software v.10.

### FluoroSpot assay

FluoroSpot assay for detecting Mouse IFN γ/IL 4 (cat# FSP-3146-10) was purchased MabTech and blocked following the manufacturer’s protocols. Then, 1 × 10^5^ isolated splenocytes were plated per well and stimulated with SIINFEKL peptide (2 μg mL^−1^ SIINFEKL) for 24 h. All tests were performed in duplicate or triplicate and included assay-positive controls. Plates were then sent to SKCCC Immune Monitoring Core for analysis.

### Enzyme-linked immunosorbent assay (ELISA)

For antibody detection, groups of C57BL/6 mice were immunized on days 0, 7, and 14. On day 30, 100 μL of blood was collected, and antigen-specific IgG levels in serum were measured by ELISA. Flat-bottom 96-well plates (Nunc) were coated overnight at 4 °C with OVA protein (2 μg per well) in 100 mM carbonate buffer (pH 9.6), then blocked with 10% fetal bovine serum (FBS) in PBS containing 0.05% Tween-20 (PBS-T). Sera were initially diluted 1:100 in PBS-T, followed by 3-fold serial dilutions. Diluted samples were added to the plates and incubated at 37 °C for 2 h. Horseradish peroxidase (HRP)-conjugated goat anti-mouse IgG (Southern Biotech, #1013-05) was used at a 1:5,000 dilution in PBS-T with 10% FBS. After incubation with HRP substrate, absorbance was measured at 450 nm using a plate reader (Thermo Fisher). A sample was considered positive if its absorbance was at least twice that of the negative control.

### Immunization and tumour therapy experiments

Mice were injected subcutaneously with 3 × 10^5^ B16-OVA cells in the therapeutic model into the right flank. Vaccinations began when tumour sizes were less than 50 mm^3^ on day 4 after tumour inoculation. Animals were immunized by i.m. injection of different LNP containing OVA mRNA as described in the main text. Tumour growth was measured every two days using a digital calliper and calculated as 0.5 × length × width × width. Mice were euthanized when the tumour volumes reached 2,000 mm^3^, and mice were also euthanized if any single tumor dimension reached 20 mm.

### nCounter Analysis System (NanoString Technology)

Total RNA was isolated from immune cells collected from muscle tissue using the Quick-RNA Microprep Kit (Zymo Research, Cat# R2062). RNA quality and concentration were assessed with a NanoDrop spectrophotometer. Gene expression profiling was performed using pre-designed NanoString nCounter CodeSets targeting mouse immune response-related genes (NanoString, Cat#115000052), according to the manufacturer’s instructions. Following hybridization, samples were processed using the nCounter Analysis System. Raw counts were normalized to internal housekeeping genes using the nSolver Analysis Software.

### Statistics & Reproducibility

Statistical analyses were performed using GraphPad Prism 8.0 and Microsoft Excel. Comparisons between two groups were conducted using two-tailed unpaired Student’s t-tests. Comparisons among more than two groups were conducted using one-way analysis of variance (ANOVA) followed by Tukey’s multiple comparisons test, with alpha set at 0.05. Survival data were analysed using the log-rank (Mantel-Cox) test, and multiple comparisons were adjusted using the Holm-Šídák method.

No statistical method was used to predetermine sample size. Sample sizes were based on prior studies, expected effect sizes, and resource availability. For vaccination studies, a minimum of n = 4 mice per group was used to permit statistical evaluation, and tumour growth studies were conducted with n = 7–8 mice per group, as stated in the figure legends. The number of biologically independent replicates is provided in each figure or figure legend.

No data were excluded from the analyses. Animal group allocation was randomized while ensuring consistent baseline characteristics. Investigators were blinded to group allocation during tumour experiments and outcome assessment. All attempts at replication were successful, and each key experiment was independently repeated at least twice with consistent results.

## Author Contributions Statement

### Author contributions

Y.Z. and H.-Q.M. conceived of and designed this study. H.-Q.M., J.P.S., and J.W.H. secured the funding for this study. Y.Z., C.W., J.M., X.L., K.D.G., X.L., D.Y., Y.S., A.H.G., L.C., and W.H.T. performed the experiments. Y.Z., C.W., and H.-Q.M. participated in data analysis and interpretation. The manuscript was written by Y.Z., C.W., and H.-Q.M., with revisions by J.W.H., J.P.S., J.C.D., and inputs from all the other authors.

## Data Availability

### Data and materials availability

All data generated or analysed during this study are provided in the paper and the Supplementary Information. No additional datasets requiring deposition in external repositories were generated.

### Code availability statement

No custom code was developed for this study.

