## Supporting Information for "Engineering age-adaptive mRNA lipid nanoparticle cancer vaccines via reprogramming systemic gene expression"

### Table of Contents

**Supplementary Figure 1 |** Gating strategy for flow cytometry plots for OVA specific T cell responses in dLNs post LNP vaccination on young and aged mice.

**Supplementary Figure 2 |** Gating strategy for flow cytometry plots for $CD8^{+}$ IFN-$\gamma^{+}$ T cell responses in spleen post LNP vaccination on young and aged mice.

**Supplementary Figure 3 |** Representative flow cytometry plots for $CD8^{+}$ IFN-$\gamma^{+}$ T cells in spleen post LNP vaccination on young and aged mice.

**Supplementary Figure 4 |** Gating strategy for flow cytometry plots for $CD3^{+}$ $CD4^{+}$ IFN-$\gamma^{+}$ T-bet$^{+}$ T cells responses in spleen post LNP vaccination on young and aged mice.

**Supplementary Figure 5 |** Gating strategy for flow cytometry plots for $CD3^{+}$ $CD4^{+}$ IL-$4^{+}$ GATA$3^{+}$ T cells responses in spleen post LNP vaccination on young and aged mice.

**Supplementary Figure 6 |** Gating strategy for flow cytometry assessment of locally recruited cells (neutrophils, DCs, macrophages and NKs) at 24h post LNP vaccination on young and aged mice.

**Supplementary Figure 7 |** Gating strategy for flow cytometry assessment of locally recruited cells (B cells and T cells) at 24h post LNP vaccination on young and aged mice.

**Supplementary Figure 8 |** Gating strategy for flow cytometry assessment of transfected cells at day 5 post LNP vaccination using Ai9 mice.

**Supplementary Figure 9 |** Gating strategy for flow cytometry assessment of antigen presentation and maturation of DCs in dLNx at day 5 post LNP vaccination.

**Supplementary Figure 10 |** Biodistribution of Cy5-labeled mRNA SM-102 LNPs in young and aged mice.

**Supplementary Figure 11 |** Gating strategy for flow cytometry assessment of Tfh cell level in spleen post LNP vaccination.

**Supplementary Figure 12 |** Gating strategy for flow cytometry assessment of CD45.1+OVA+ T cells in spleen.

**Supplementary Figure 13 |** Biodistribution of Cy5-labeled mRNA B LNPs in young and aged mice.

**Supplementary Figure 14 |** Serum OVA-specific IgG1 and IgG2c responses in young and aged mice following intramuscular vaccination with B LNPs.

**Supplementary Figure 15 |** Analysis of tumor-infiltrating lymphocytes on day 14 post-tumor inoculation following treatment with B LNPs in young and aged mice.

**Supplementary Table 1 |** Composition details and characterization of the six evaluated LNP formulations.

**Supplementary Table 2 |** Anti-mouse antibodies used in flow cytometry panels. Marker, fluorophore, catalogue number, source, and concentration are indicated.

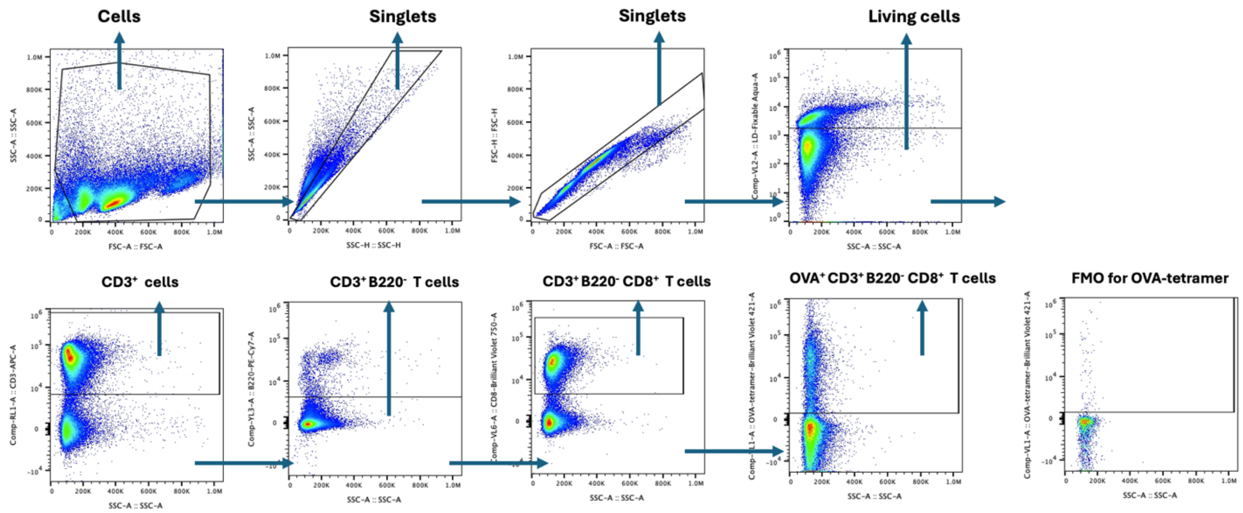


**Supplementary Figure 1. Gating strategy for flow cytometry plots for OVA specific T cell responses in dLNs post LNP vaccination on young and aged mice.** Initially, lymphocytes isolated from the dLNs were selected using SSC-A and FSC-A parameters, followed by singlet selections with SSC-A/SSC-H and FSC-A/FSC-H plots. Viable cells were identified and selected based on the live/dead Fixable Aqua-A and SSC-A plot. Next, the $CD3^{+}B220^{-}CD8^{+}$ T cell population was selected with downstream analysis focused on antigen (OVA)-specific and $CD8^{+}$ T cell subtypes.

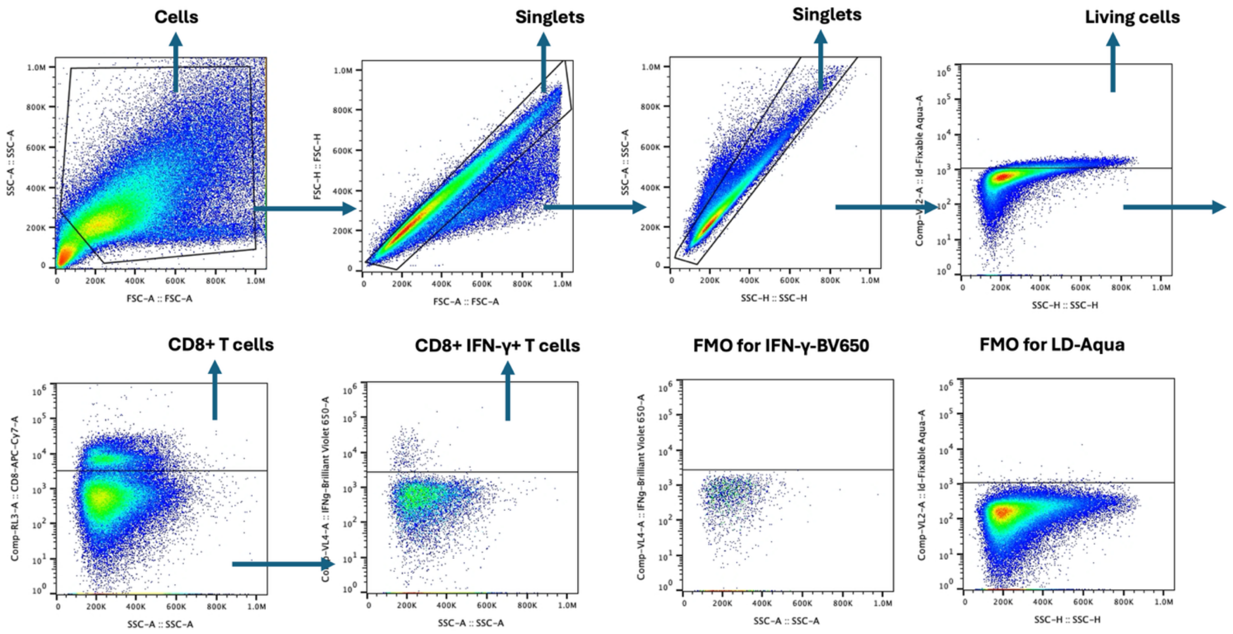


**Supplementary Figure 2. Gating strategy for flow cytometry plots for CD8+ IFN-γ+ T cell responses in spleen post LNP vaccination on young and aged mice.** Initially, lymphocytes isolated from the spleen were selected using SSC-A and FSC-A parameters, followed by singlet selections with FSC-A/FSC-H and SSC-A/SSC-H plots. Viable cells were identified and selected based on the live/dead Fixable Aqua-A and SSC-A plot. Next, the $CD8^+$ T cell population was selected with downstream analysis focused on IFN-$\gamma^+$ T cell subtypes.

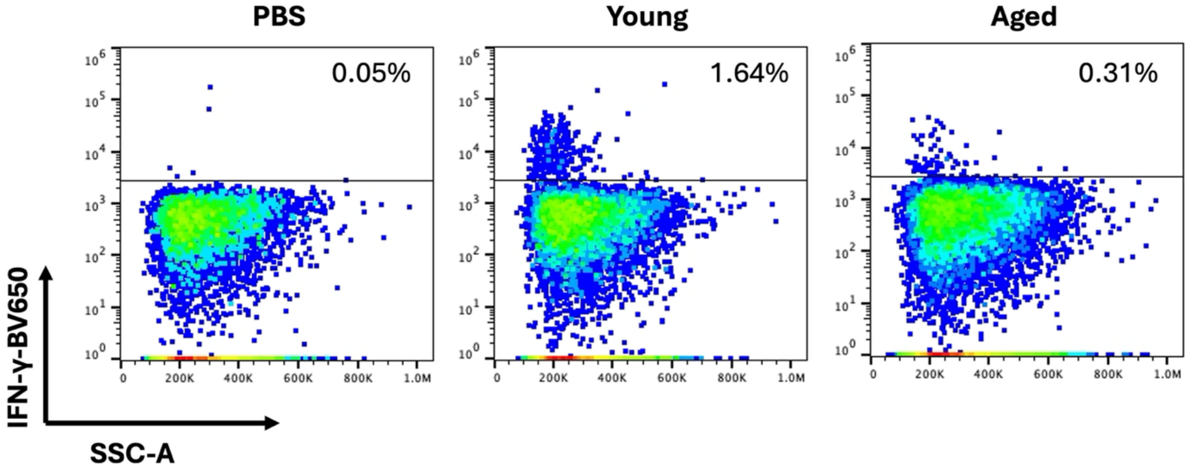


**Supplementary Figure 3. Representative flow cytometry plots for CD8+ IFN-γ+ T cells in spleen post LNP vaccination on young and aged mice.** Lymphocytes isolated from the spleen were restimulated *in vitro* with SIINFEKL peptide (1 µg/mL SIINFEKL) for 12 h and assessed via intracellular cytokine staining and flow cytometry and to determine the percentages of $CD8^{+}$ IFN-$\gamma^{+}$ T cells.

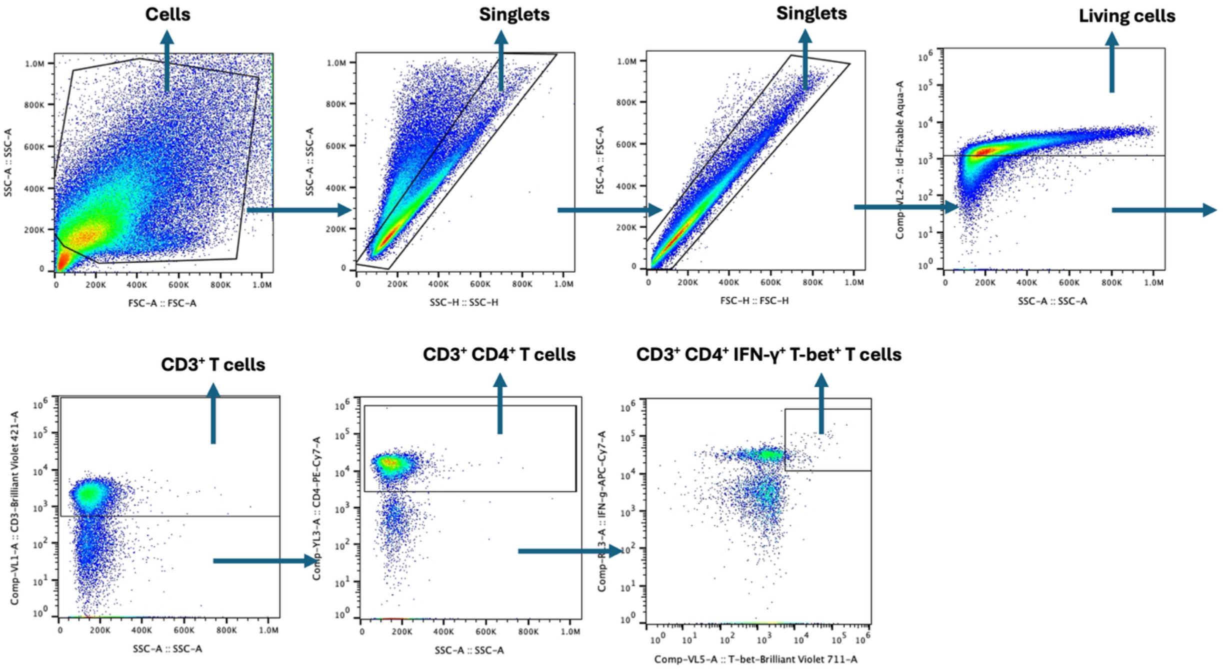


**Supplementary Figure 4. Gating strategy for flow cytometry plots for CD3$^+$ CD4$^+$ IFN-γ$^+$ T-bet$^+$ T cells responses in spleen post LNP vaccination on young and aged mice.** Lymphocytes isolated from the spleen were restimulated *in vitro* with SIINFEKL peptide (1 µg/mL SIINFEKL) for 12 h and assessed via intracellular cytokine staining and flow cytometry and to determine the percentages of CD3$^+$ CD4$^+$ IFN-γ$^+$ T-bet$^+$ T cells. Initially, lymphocytes isolated from the spleen were selected using SSC-A and FSC-A parameters, followed by singlet selections with SSC-A/SSC-H and FSC-A/FSC-H plots. Viable cells were identified and selected based on the live/dead Fixable Aqua-A and SSC-A plot. Next, the CD3$^+$ CD8$^+$ T cell population was selected with downstream analysis focused on T-bet$^+$ IFN-γ$^+$ T cell subtypes.

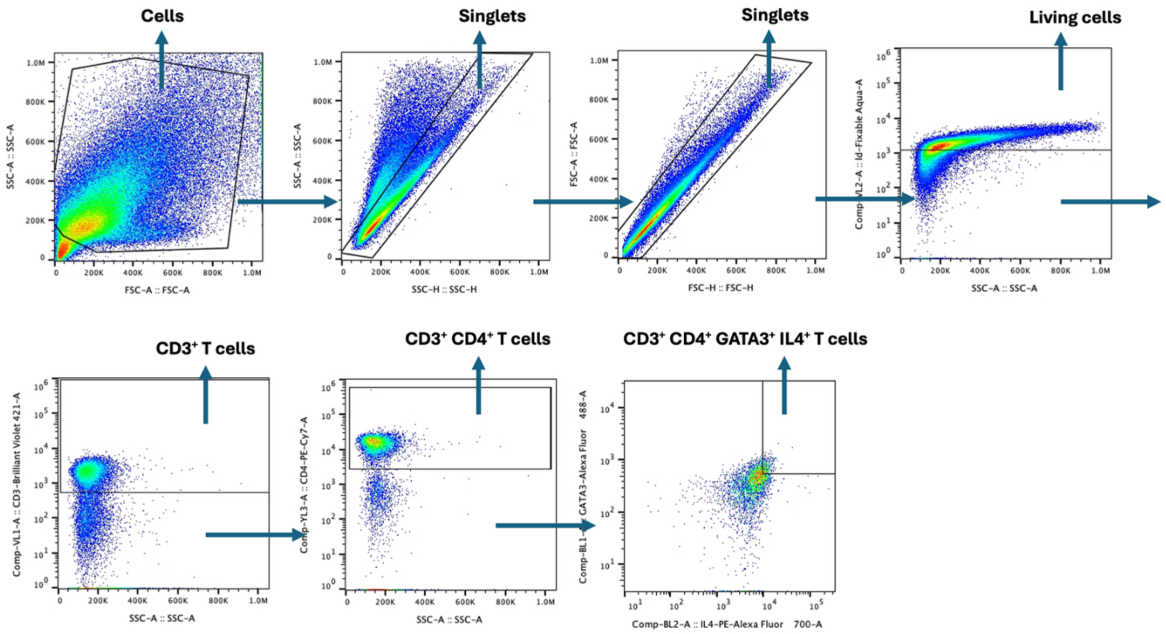


**Supplementary Figure 5. Gating strategy for flow cytometry plots for CD3$^+$ CD4$^+$ IL-4$^+$ GATA3$^+$ T cells responses in spleen post LNP vaccination on young and aged mice.** Lymphocytes isolated from the spleen were restimulated *in vitro* with SIINFEKL peptide (1 µg/mL SIINFEKL) for 12 h and assessed via intracellular cytokine staining and flow cytometry and to determine the percentages of CD3$^+$ CD4$^+$ IL-4$^+$ GATA3$^+$ T cells. Initially, lymphocytes isolated from the spleen were selected using SSC-A and FSC-A parameters, followed by singlet selections with SSC-A/SSC-H and FSC-A/FSC-H plots. Viable cells were identified and selected based on the live/dead Fixable Aqua-A and SSC-A plot. Next, the CD3$^+$ CD8$^+$ T cell population was selected with downstream analysis focused on IL-4$^+$ GATA3$^+$ T cell subtypes.

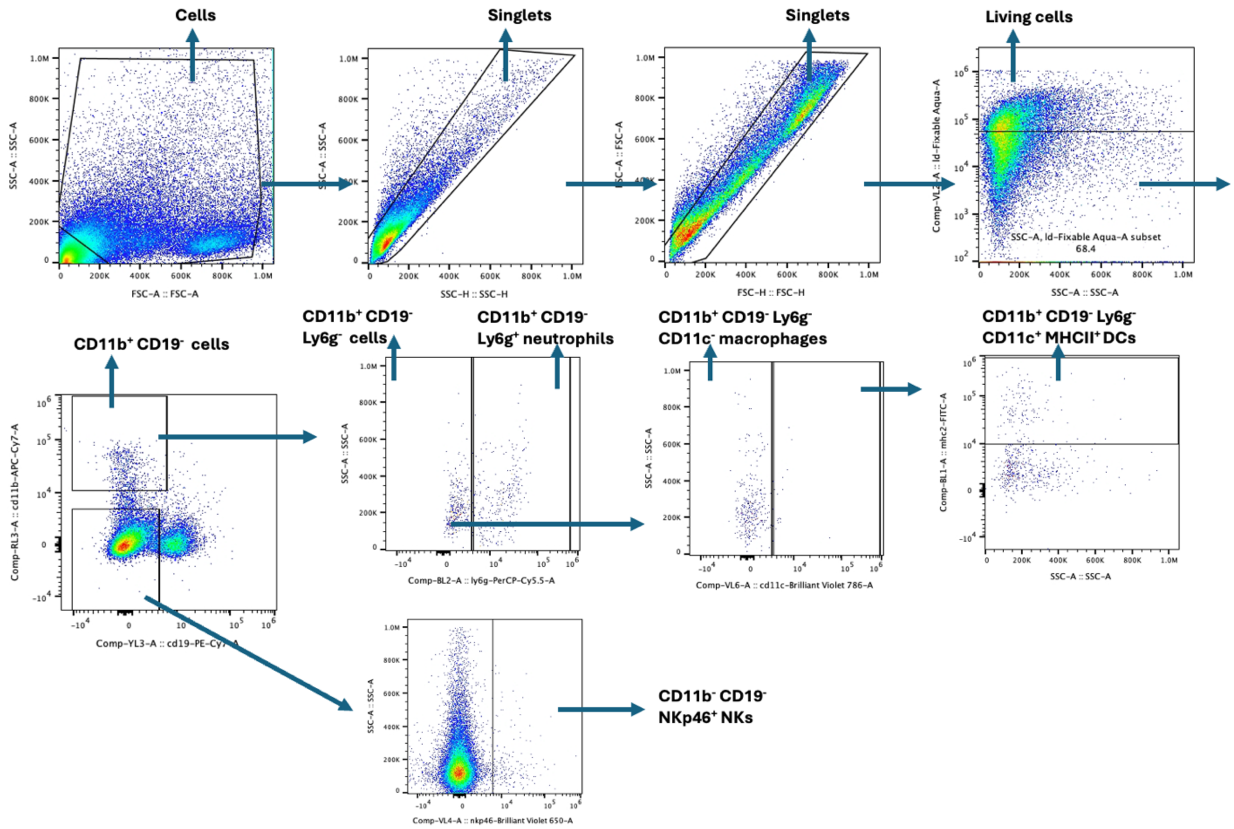


**Supplementary Figure 6. Gating strategy for flow cytometry assessment of locally recruited cells (neutrophils, DCs, macrophages and NKs) at 24h post LNP vaccination on young and aged mice.** Initially, cells isolated from the muscle injection site were selected using SSC-A and FSC-A parameters, followed by singlet selections with SSC-A/SSC-H and FSC-A/FSC-H plots. Viable cells were identified and selected based on the live/dead Fixable Aqua-A and SSC-A plot. Next, $CD19^{-}CD11b^{+}$ cells were gated, and Ly6g+ cells were further characterized as neutrophils. Additionally, $Ly6g^{-}CD11c^{+}$ $MHCII^{+}$ cells were identified as dendritic cells (DCs), while $Ly6g^{-}CD11c^{-}$ cells were designated as macrophages. $CD19^{-}CD11b^{-}NKp46^{+}$ cells were identified as NK cells.

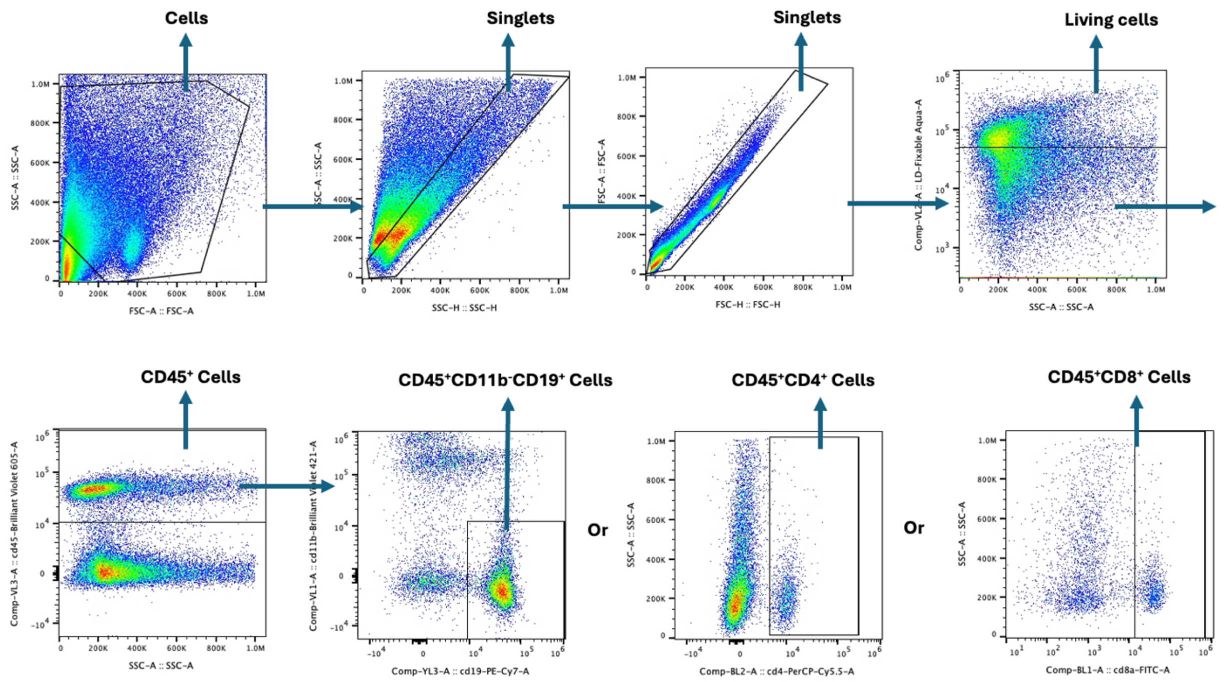


**Supplementary Figure 7. Gating strategy for flow cytometry assessment of locally recruited cells (B and T cells) at 24h post LNP vaccination on young and aged mice.** Initially, cells isolated from the muscle injection site were selected using SSC-A and FSC-A parameters, followed by singlet selections with SSC-A/SSC-H and FSC-A/FSC-H plots. Viable cells were identified and selected based on the live/dead Fixable Aqua-A and SSC-A plot. Next, $CD45^+$ cells were gated, and $CD11b^-CD19^+$ cells were further characterized as B cells. Additionally, $CD45^+CD4^+$ cells were identified as CD4 T cells, while $CD45^+CD8^+$ cells were designated as CD8 T cells.

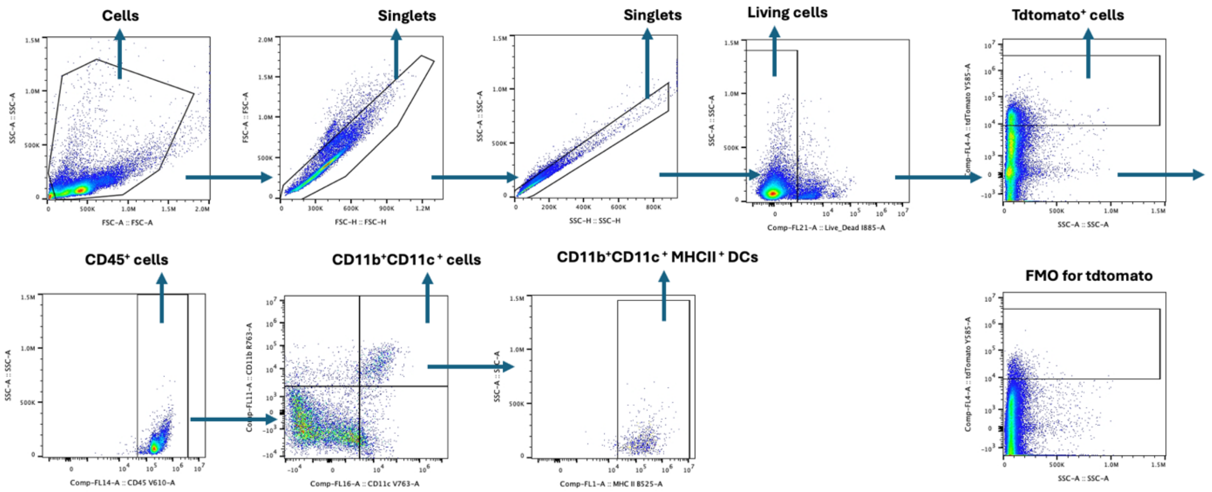


**Supplementary Figure 8. Gating strategy for flow cytometry assessment of transfected cells at day 5 post LNP vaccination using Ai9 mice.** Initially, cells isolated from the tissue including muscle injection site and dLNs were selected using SSC-A and FSC-A parameters, followed by singlet selections with SSC-A/SSC-H and FSC-A/FSC-H plots. Viable cells were identified and selected based on the live/dead Fixable Aqua-A and SSC-A plot. Next, CD45$^+$ cells were gated, and CD11b$^+$CD11c$^+$ MHCII$^+$ cells were further characterized as DC cells. Additionally, CD45$^+$CD11b$^+$ CD11c$^-$ cells were identified as macrophages.

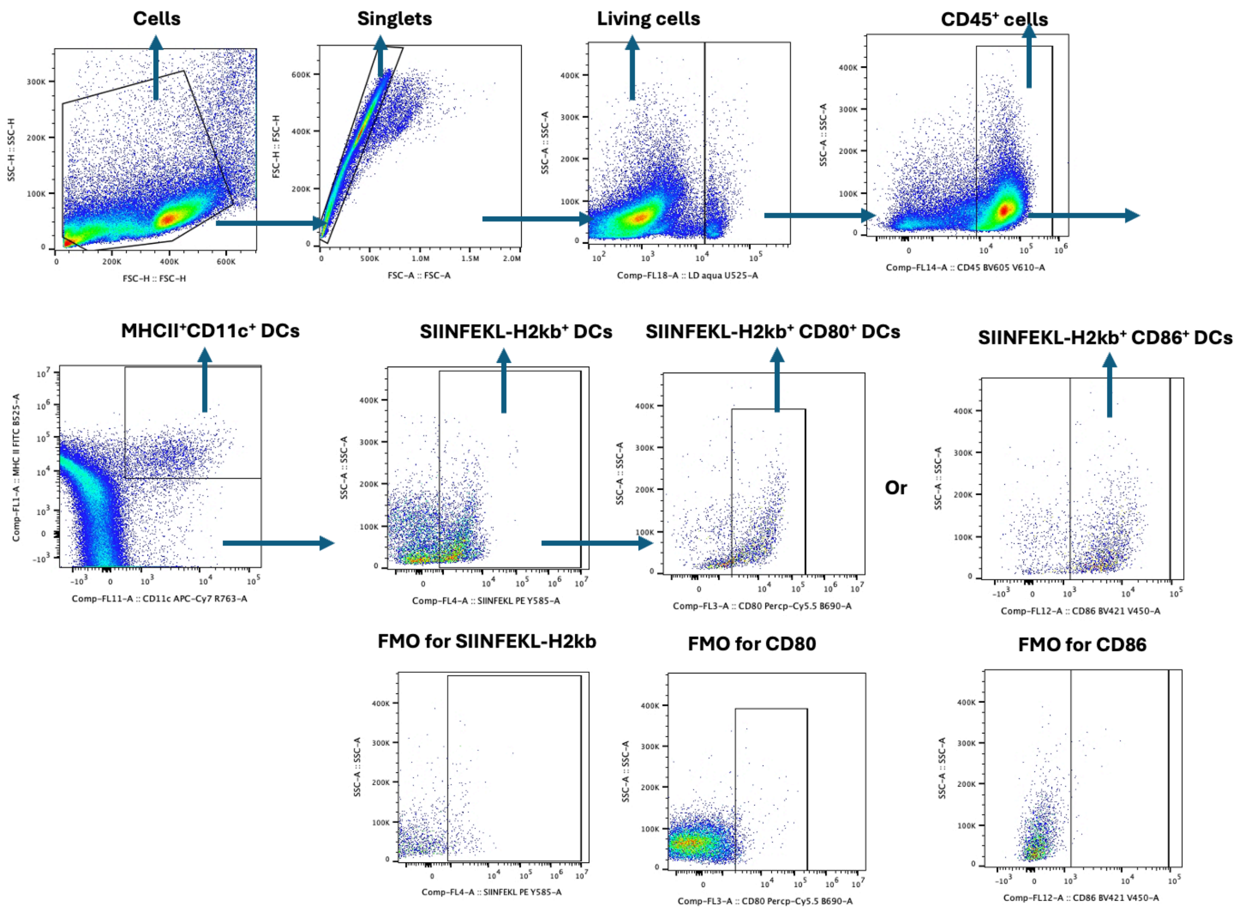


**Supplementary Figure 9. Gating strategy for flow cytometry assessment of antigen presentation and maturation of DCs in dLNx at day 5 post LNP vaccination.** Initially, cells isolated from the dLNs were selected using SSC-H and FSC-H parameters, followed by singlet selections with FSC-A/FSC-H plots. Viable cells were identified and selected based on the live/dead Fixable Aqua-A and SSC-A plot. Next, MHCII$^+$ CD11c$^+$ cells were gated, were characterized as DC cells, and SIINFEKL-H2Kb$^+$ level were meassured. Additionally, CD80$^+$ or CD86$^+$ cells were identified as matured DCs.

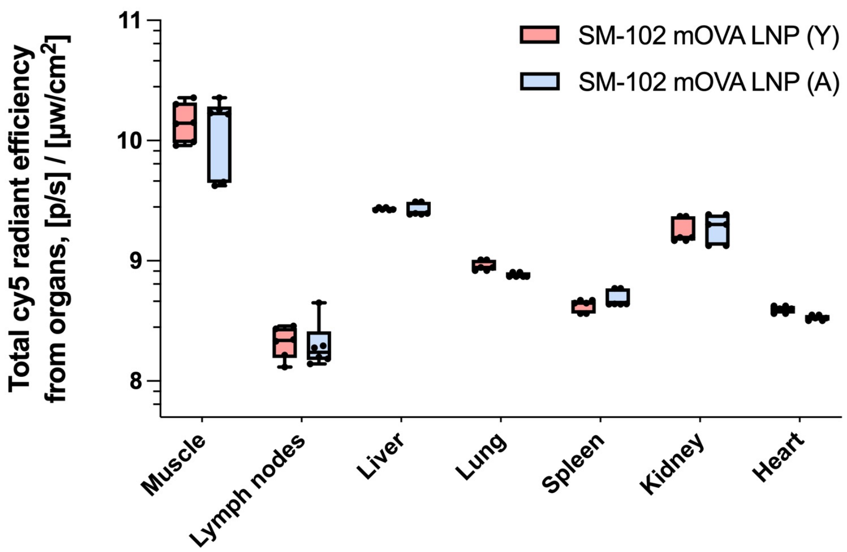


**Supplementary Figure 10. Biodistribution of Cy5-labeled mRNA SM-102 LNPs in young and aged mice.** IVIS imaging of major organs 24 hours after intramuscular injection of Cy5-labeled mRNA formulated in LNPs (10 µg per mouse) in young (6–8 weeks) and aged (10–12 months) C57BL/6 mice. Organs assessed include liver, lungs, spleen, kidneys, heart, injection-site muscle, and dLNs. Fluorescence intensity reflects mRNA biodistribution and retention across tissues.

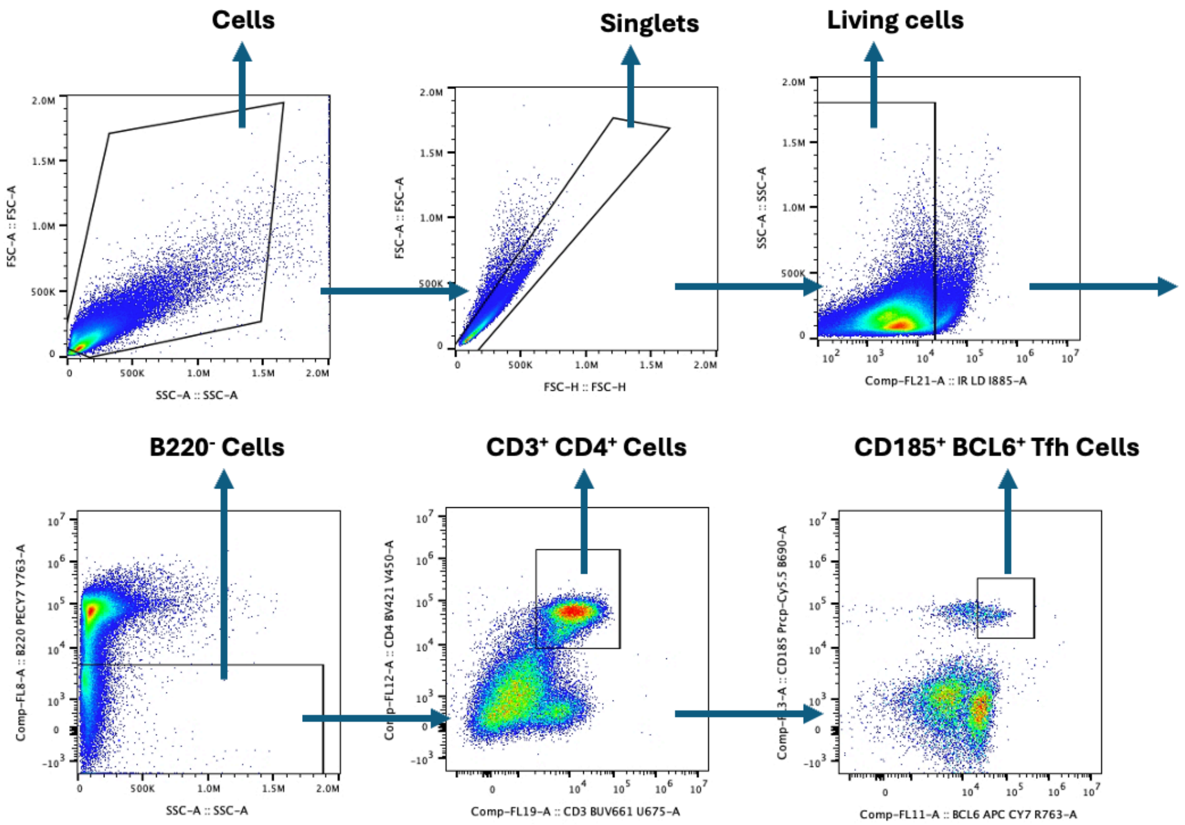


**Supplementary Figure 11. Gating strategy for flow cytometry assessment of Tfh cell level in spleen post LNP vaccination.** Lymphocytes isolated from the spleen were restimulated *in vitro* with SIINFEKL peptide (1 μg/mL SIINFEKL) for 12 h and assessed via intracellular cytokine staining and flow cytometry and to determine the percentages of B220$^-$ CD3$^+$ CD4$^+$ BCL-6$^+$ CD185$^+$ Tfh cells. Initially, lymphocytes isolated from the spleen were selected using SSC-A and FSC-A parameters, singlets were gated through FSC-A and FSC-H. Viable cells were identified and selected based on the live/dead Fixable IR-A and SSC-A plot. Next, B220$^-$ CD3$^+$ CD4$^+$ cells were gated, were characterized as CD4 T cells, and CD185$^+$ BCL6$^+$ cells were identified as Tfh cells.

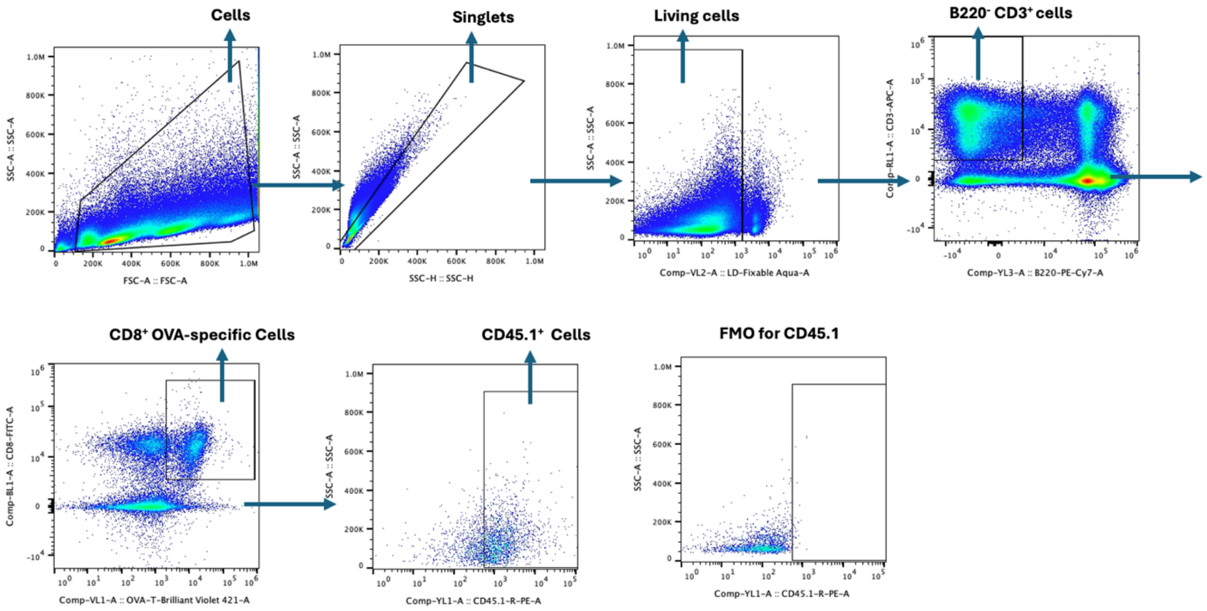


**Supplementary Figure 12. Gating strategy for flow cytometry assessment of CD45.1$^+$OVA$^+$ T cells in spleen.** Initially, lymphocytes isolated from the spleen were selected using SSC-A and FSC-A parameters, singlets were gated through FSC-A and FSC-H. Viable cells were identified and selected based on the live/dead Fixable IR-A and SSC-A plot. Next, B220$^-$ CD3$^+$ CD8$^+$ OVA$^+$ cells were gated, and CD45.1$^+$ cells were identified.

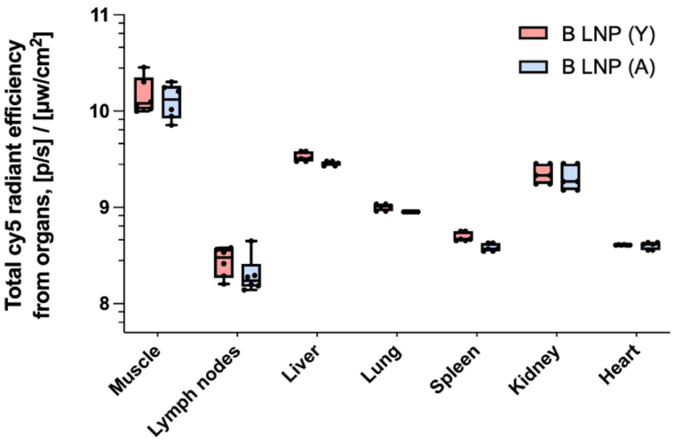


**Supplementary Figure 13. Biodistribution of Cy5-labeled mRNA B LNPs in young and aged mice.** IVIS imaging of major organs 24 hours after intramuscular injection of Cy5-labeled mRNA formulated in B LNPs (10 µg per mouse) in young (6–8 weeks) and aged (10–12 months) C57BL/6 mice. Organs assessed include liver, lungs, spleen, kidneys, heart, injection-site muscle, and dLNs. Fluorescence intensity reflects mRNA biodistribution and retention across tissues.

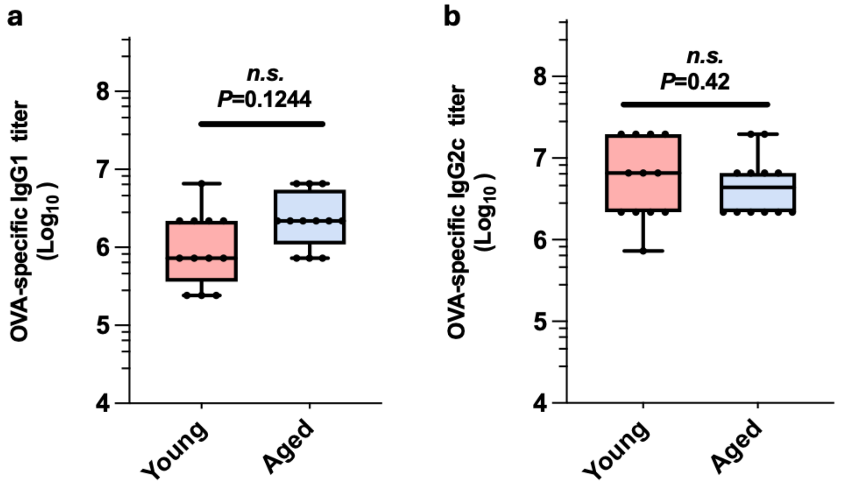


**Supplementary Figure 14. Serum OVA-specific IgG1 and IgG2c responses in young and aged mice following intramuscular vaccination with B LNPs.** Young (6–8 weeks) and aged (10–12 months) C57BL/6 mice were vaccinated i.m. with 10 µg mOVA-loaded B LNPs on days 0, 7, and 14, and analyzed on day 30. Serum OVA-specific IgG1 (**a**) IgG2c (**b**) titers were measured by ELISA (n=12 mice). Data were analyzed using unpaired t-test. NS, not significant.

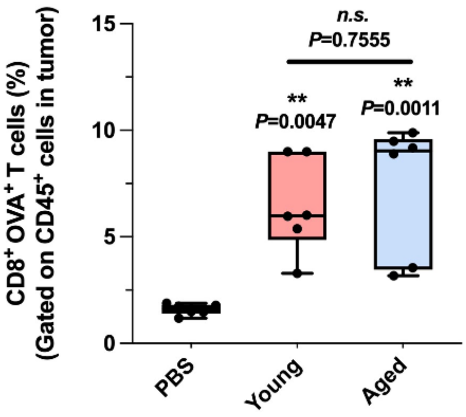


**Supplementary Figure 15. Analysis of tumor-infiltrating lymphocytes on day 14 post-tumor inoculation following treatment with B LNPs in young and aged mice.** Mice were subcutaneously inoculated with B16-OVA tumor cells on day 0, followed by mRNA-LNP vaccination (10 μg B mOVA LNP, i.m.) on days 4, 11, and 18. Tumors were dissociated into single-cell suspensions and analyzed by flow cytometry. Shown are the frequency of $CD45^{+}CD8^{+}OVA^{+}$ T cells among total live cells. Data were analyzed using two-way ANOVA and Tukey's multiple comparisons test. **P < 0.01; NS, not significant.

**Supplementary Table 1. Composition details and characterization of the six evaluated LNP formulations.**

| Formulation Code | A LNP | B LNP | C LNP | D LNP | E LNP | SM-102 LNP |
|---|---|---|---|---|---|---|
| *Composition (molar ratio):* | | | | | | |
| Ionizable lipid | 49.74 (Dlin-MC3-DMA) | 46.3 (ALC-0315) | 40 (Dlin-MC3-DMA) | 36.36 (Dlin-MC3-DMA) | 54.55 (Dlin-MC3-DMA) | 50 (SM-102) |
| Helper lipid | 10.26 (DSPC) | 9.4 (DSPC) | 40 (DOPE) | 3.64 (DSPC) | 5.45 (18 PG) | 10 (DSPC) |
| Cholesterol | 38.46 | 42.7 | 19.96 | 59.88 | 39.92 | 38.5 |
| DMG-PEG2000 | 1.54 | 1.6 | 0.04 | 0.12 | 0.08 | 1.5 |
| N/P ratio | 4 | 6 | 4 | 8 | 8 | 6 |
| *Formulation features:* | | | | | | |
| Z-average diameter (nm) | 126.6 ± 0.9 | 126.4 ± 0.6 | 162.9 ± 2.6 | 125.6 ± 0.6 | 139.1+4.0 | 117.6 ± 1.0 |
| Average PDI | 0.208 ± 0.022 | 0.212 ± 0.018 | 0.209 ± 0.009 | 0.181 ± 0.012 | 0.209 ± 0.005 | 0.115 ± 0.028 |
| Average Zeta potential (mV) | -2.95 ± 0.51 | -2.92 ± 0.20 | -11.0 ± 2.30 | -1.25 ± 0.50 | -5.83 ± 1.37 | -2.80 ± 0.71 |
| Average EE% | 99.95 | 96.45 | 99.03 | 99.96 | 99.92 | 99.85 |

**Supplementary Table 2. Anti-mouse antibodies used in flow cytometry panels. Marker, fluorophore, catalogue number, source, and concentration are indicated.**

| Antigen | Fluorophore | Catalogue # | Source | Concentration |
|---|---|---|---|---|
| CD8a | APC-Cy7 | 100714 | Biolegend | 1:200 dilution |
| OVA SIIGFEKL (Tetramer) | Brilliant Violet 421 | N/A | NIH tetramer core facility | 1:400 dilution |
| CD3 | APC | 100236 | Biolegend | 1:100 dilution |
| Live/Dead | Live/Dead Fix Aqua | L34957 | Thermo Fisher | 1:1000 dilution |
| CD45 | FITC | 103108 | Biolegend | 1:250 dilution |
| CD19 | PE-Cy7 | 25019382 | Thermo Fisher | 1:200 dilution |
| Ly6G | PerCP/Cy5.5 | 127616 | Biolegend | 1:150 dilution |
| CD45 | Brilliant Violet 421 | 103134 | Biolegend | 1:250 dilution |
| CD3 | PE | 100206 | Biolegend | 1:200 dilution |
| CD8 | FITC | 100706 | Biolegend | 1:200 dilution |
| Fixable Live/Dead | Live/Dead Fix Near IR (780) | L10119 | Thermo Fisher | 1:1000 dilution |
| CD8a | Brilliant Violet 750 | 747134 | Biolegend | 1:200 dilution |
| CD45R | PE-Cy7 | 103222 | Biolegend | 1:100 dilution |
| NKp46 | BV650 | 137635 | Biolegend | 1:100 dilution |
| TNF-a | Alexa Fluor 700 | 506338 | Biolegend | 1:100 dilution |
| CD3 | Brilliant Violet 421 | 100228 | Biolegend | 1:200 dilution |
| IFN-g | BV650 | 505832 | Biolegend | 1:50 dilution |
| IL-4 | PERCP-eFluor 710 | 46704182 | Thermo Fisher | 1:50 dilution |
| IFN-g | APC/CY7 | 505850 | Biolegend | 1:200 dilution |
| GATA3 | Alexa Fluor 488 | 653808 | Biolegend | 1:50 dilution |
| T-bet | BV711 | 644820 | Biolegend | 1:50 dilution |
| CD45.1 | PE | 110708 | Biolegend | 1:100 dilution |
| Bcl-6 | APC-Cy7 | 563581 | BD | 1:50 dilution |
| CXCR5 (CD185) | PerCP/Cy5.5 | 145508 | Biolegend | 1:100 dilution |
| MHCII | FITC | 107606 | Biolegend | 1:200 dilution |

| | | | | |
|---|---|---|---|---|
| CD11c | BV750 | 117357 | Biolegend | 1:200 dilution |
| H2KB-SIINFEKL | PE | 12574382 | Thermo Fisher | 1:200 dilution |
| CD80 | PerCP/Cy5.5 | 104722 | Biolegend | 1:100 dilution |
| CD86 | BV421 | 105123 | Biolegend | 1:100 dilution |
| CD4 | PE/CY7 | 100422 | Biolegend | 1:100 dilution |
